# Motor nerves direct the development of the sympathetic nervous system

**DOI:** 10.1101/2023.03.17.533145

**Authors:** Alek Erickson, Alessia Motta, Maria Eleni Kastriti, Steven Edwards, Francois Lallemend, Saida Hadjab, Christiana Ruhrberg, Quenten Schwarz, Kaj Fried, Dario Bonanomi, Igor Adameyko

**Affiliations:** Department of Physiology and Pharmacology, Karolinska Institutet, Stockholm, 17165, Sweden; Division of Neuroscience, San Raffaele Hospital, 20132 Milano, Italy; Center for Brain Research, Medical University Vienna, Vienna, 1090, Austria; Department of Applied Physics, KTH Royal Institute of Technology, Stockholm, 10044, Sweden; Department of Neuroscience, Biomedicum, Karolinska Institute, Stockholm, 17165, Sweden; University College London, Department of Ophthalmology London, United Kingdom; Center for Cancer Biology, University of South Australia, Adelaide, Australia

## Abstract

The sympathetic nervous system controls a wide spectrum of bodily functions including operation of vessels, cardiac rhythm, and the “flight or fight response”. Sympathetic neurons, which are neural crest-derived, develop in coordination with presynaptic motor nerves extending from the central nervous system (CNS). By using nerve-selective genetic ablations, we revealed that sympathetic ganglia development depends on CNS-derived motor innervation. In the absence of preganglionic motor nerves, trunk sympathetic chain ganglia were fragmented and smaller in size, while cervical ganglia were severely misshapen. Sympathetic neurons were misplaced along sensory fibers and projected towards abnormal paths, in some cases invading the sensory dorsal root ganglia. The misplaced progenitors of sympathoblasts corresponded to the nerve-associated, neural crest-derived Schwann cell precursors (SCPs). Notably, we found that SCPs activate the autonomic marker PHOX2B while migrating along motor nerves towards the region of the dorsal aorta in wildtype embryos, suggesting that SCP differentiate into sympathetic neurons while still nerve-associated in motor-ablated embryos. Ligand-receptor prediction from single cell transcriptomic data coupled with functional studies identified Semaphorin 3A/3F as candidate motor nerve-derived signals influencing neural crest migration along axons. Thus, motor nerves control the placement of sympathoblasts and their subsequent axonal navigation during critical periods of sympathetic chain development.

## INTRODUCTION

The major physiological functions within our bodies, including gut motility, heart rate, saliva production, thermal control and stress responses are regulated by fine nerves belonging to the autonomic compartment of the peripheral nervous system (PNS) ^1^. There are multiple subdivisions of the autonomic nervous system (ANS) including the enteric branch and the rather antagonistic sympathetic and parasympathetic branches. The sympathetic branch is essential for the “fight of flight” response, and largely consists of paired ganglia segmented along the anteroposterior axis, lateral to the dorsal aorta. These ganglia are innervated by the visceral cholinergic motor nerves originating from the ventral neural tube ^2^. The development of these motoneurons requires the expression of the transcriptional factor *Olig2* for their maturation ^3,4^. Another main branch, composed of parasympathetic neurons, antagonizes the sympathetic branch in the majority of locations to control the “rest and digest” response ^5^. The proper functionality, growth, and development of ANS branches are each essential for homeostasis and survival of all mammals.

All neurons and glial cell types of the autonomic nervous system are derived from the transient, migratory embryonic cell population called the neural crest ^6^. Some develop as a result of the immediate neural crest migration and differentiation (for instance, the majority of sympathetic populations and some enteric neurons), and some develop later via the intermediate stage represented by the neural crest-derived and nerve-associated Schwann cell precursors (SCPs). For instance, parasympathetic ganglia ^7 8^ and chromaffin cells of adrenal and extra-adrenal organs ^9 10^ as well as some parts of the enteric nervous system ^11^ originate from SCPs associated with preganglionic nerve fibers.

In the trunk, neural crest cells form within the dorsal neural tube during neural tube closure, and their migration begins around embryonic day 9 in the mouse embryo ^12 13^. The early streams of migrating neural crest cells proceed towards the dorsal aorta, where extrinsic signals instruct differentiation of neural crest into coalescing sympathetic ganglia ^14^. At the same time, numerous neural crest cells settle on the outgrowing motor and sensory nerves and become SCPs – another multipotent cell type highly similar to their mother population of the neural crest^15^. SCPs spread by migrating along the outgrowing nerves and reach all innervated locations, where some of them detach and give rise to parasympathetic ganglia, certain populations of sympathetic and enteric neurons, and nearly all chromaffin cells. Though the majority of the neurons within the sympathetic chain is thought to form from the early neural crest streams^16^, disentangling the co-contribution of primeval SCPs to sympathetic ganglia has been challenging because of the rapid and highly parallel development of the peripheral nervous system during organogenesis.

Studies characterizing the phenotype of mouse embryos with specific nerve ablations have revealed that various aspects of nervous system development depend on the correct positioning and growth of pre-existing nerve tracts ^17,18^. This general wiring strategy often involves the initial outgrowth of “pioneering” neuronal populations that establish a template guiding the axons of follower neurons ^19 20 21^. This is evident in the pathfinding of some trunk sympathetic and sensory fibers that rely on pre-extended spinal motor neuron projections^18^. In addition, the formation of many parasympathetic ganglia strictly depends on motor or sensory innervation to supply progenitor cells in the form of SCPs. However, in the case of sympathetic ganglia, which receive direct contribution from early neural crest, innervation is dispensable for the early induction and overall positioning of the sympathetic chain. By extension, motor and viscerosensory innervation has also been considered unnecessary for sympathetic ganglia formation ^22^. For these reasons, the attribute of “nerve-independence” is now thought to be associated with development of the sympathetic branch, and has been used as a key criterion in recent debates on whether the sacral outflow can be classified as truly parasympathetic or sympathetic ^23 24^. Notwithstanding, it has been recently discovered that some sympathetic neurons in the paraganglia, as well as intra-adrenal sympathetic neurons, are derived from visceral motor nerve-associated SCPs ^25 10^

In this study, we addressed the question of how visceral motor nerves contribute to development of the sympathetic ganglia. We discovered that after genetic ablation of motor nerves, the sympathetic chain becomes fragmented and surrounded by small ectopic clusters of sympathetic neurons that often extend axons into the dorsal root ganglia. These misplaced sympathetic cells are likely derived from sensory-associated SCPs and satellite glial cells, as they are invariably associated with sensory neurons, and project along them. The results suggest motor nerves affect the positioning of sympathetic neuroblasts and influence sympathetic axonal navigation.

## RESULTS

### Sympathetic ganglia defects in motor nerve-ablated mouse embryos

To investigate the role of motor nerves in mouse sympathetic ganglia development, we utilized *Olig2*^*Cre*^; *R26R*^*DTA*^ and *Hb9*^*Cre*^*;Isl2*^*DTA*^ transgenic mouse models in which motor neurons are selectively ablated following the expression of diphtheria toxin (DTA) in motor neuron progenitors with *Olig2*^*Cre*^, or in postmitotic motor neurons based on overlapping expression of *Hb9* and *Isl2* ^18,26-28^. *Olig2*^*Cre*^; *R26R*^*DTA*^ embryos displayed a near total loss of HB9^+^ motor neurons by E10.5 (Supplementary Figure 1A) and E11.5 (Supplementary Figure 1B). The motor-less embryos showed no significant difference in body length at E12.5 (Supplementary Figure 2A) or in the size of the E10.5 sympathetic anlagen. (Supplementary Figure 2B). Likewise, there was not any obvious changes in the cross-sectional diameter of the tightly condensed regions of forming sympathetic ganglia at E11.5 (Supplementary Figure 2C). Together, this suggests that overall embryonic growth, as well as early waves of neural crest migration, are not affected in these embryos. We also compared the *Olig2*^*Cre*^; *R26R*^*DTA*^ embryos to previously reported transgenic models of motor ablation. For instance, studies of *Olig2* knockout mice report pelvic ganglion development was unaffected ^22^, and the same was observed in *Olig2*^*Cre*^; *R26R*^*DTA*^ embryos (Supplementary Figure 2D and 2E). Furthermore, *Hb9*^*Cre*^*;Isl2*^*DTA*^ embryos were previously reported to have reduced numbers of chromaffin cells in the adrenal medulla ^26^, and this was also observed in *Olig2*^*Cre*^; *R26R*^*DTA*^ embryos (Supplementary Figure 2F), suggesting this transgenic model was an effective and specific way to eliminate motor neurons from the early embryo.

To examine how sympathetic ganglia develop in the absence of motor nerves in *Olig2*^*Cre*^; *R26R*^*DTA*^ embryos we used whole mount immunostaining to visualize expression of Tyrosine Hydroxylase (TH) and Neurofilament 200kD (hereafter referred to as 2H3) with light sheet microscopy. At E11.5, the sympathetic chain in *Olig2*^*Cre*^; *R26R*^*DTA*^ embryos, but not control embryos, was surrounded by a constellation of ectopic TH^+^ cells along the entire trunk (Figure 1A), near 2H3^+^ nerve fibers. Defects in the compaction of sympathetic ganglia was also found at E12.5 (Figure 1B, middle column) in mutant but not control embryos, and by this point in development also begin to project abnormally (Figure 1B, right column). In the lumbosacral region, in addition to the ectopic sympathoblasts, chain ganglia formed discrete aberrant clusters, unlike the regular and continuous posterior sympathetic chain of control littermates (Figure 1B, lower panels). This striking “fragmented” phenotype, characterized by ectopic, misplaced clusters of sympathetic neurons was observed at multiple stages (E11.5, E12.5, E13.5, E18.5) in *Olig2*^*Cre*^; *R26R*^*DTA*^ embryos but not in control littermates, suggesting that the defect is not rooted in an abnormal developmental schedule (Figure 1A-B, Figure 2, Figure 5). Quantification of contiguous TH^+^ volumes of *Olig2*^*Cre*^; *R26R*^*DTA*^ embryo and control littermates at indicated that sympathetic ganglia were similar sizes at E11.5 (Figure 1C) but that at E12.5, the TH^+^ volume was significantly smaller in the mutant embryos, compared to control (Figure 1D). Specifically, post-cervical sympathetic chains, as well as adrenal tissues, were significantly smaller in the mutants, while the volume of the cervical ganglia, which did not appear fragmented, was not significantly smaller (Figure 1E). Thus, loss of motor nerves in *Olig2*^*Cre*^; *R26R*^*DTA*^ embryos leads to misplaced sympathoblasts and ganglia fragmentation, overall resulting in smaller sympathetic chains. Interestingly, misplaced sympathetic neurons grew abnormal projections that extended up to millimeters farther than in controls (Figure 1F) resulting in improper innervation of the dorsal root ganglia (DRGs) and limbs (Figure 1B, right panels). This data suggests that motor neurons might regulate sympathetic axon growth, either directly or as a consequence of influencing the positioning of trunk sympathetic neurons.

**Figure 1:**
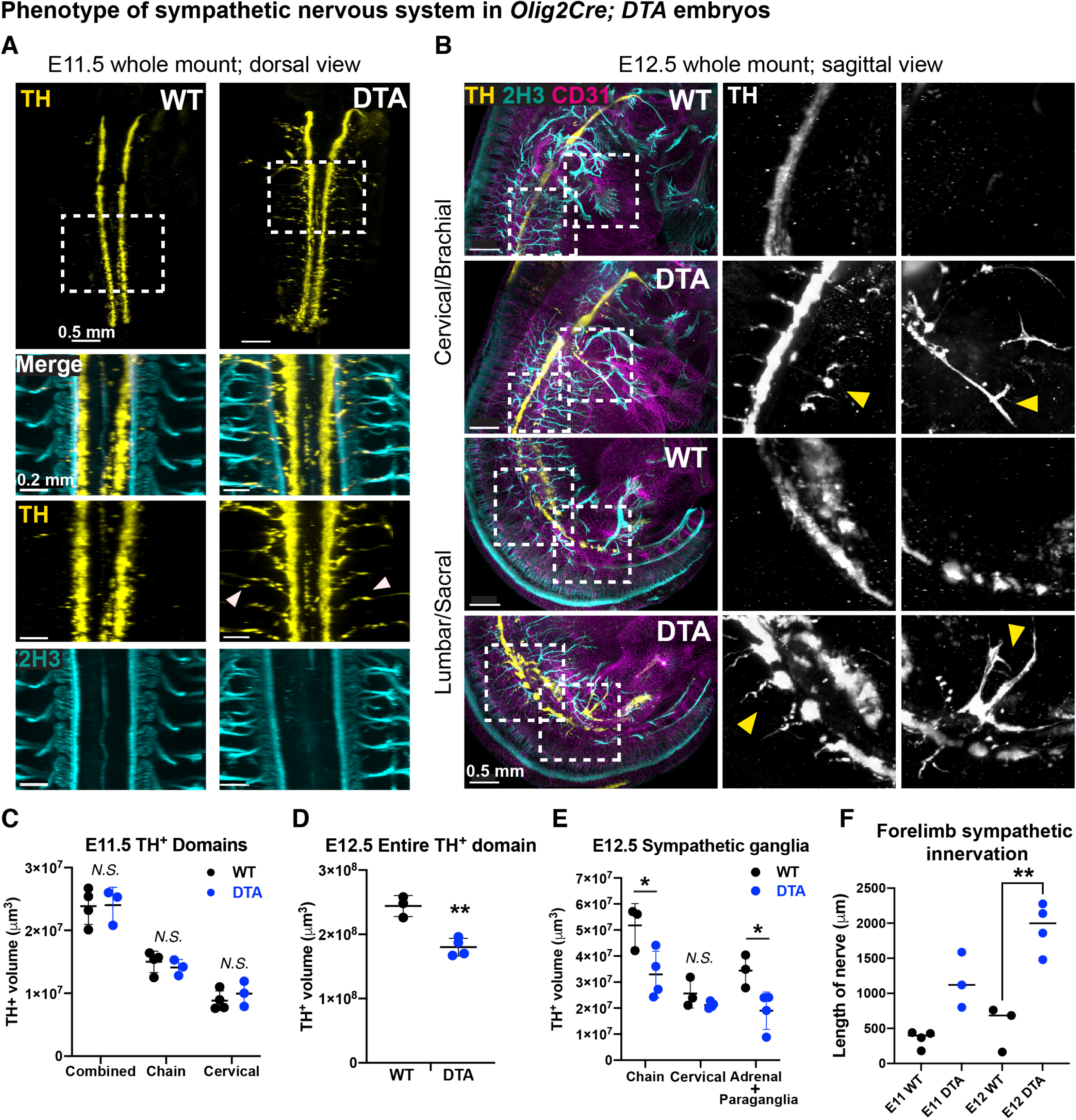
Sympathetic chain ganglia development depends on visceromotor innervation. **(A)** Dorsal view of a whole mount immunostaining for TH and Neurofilament 200kD (2H3) of E11.5 *Olig2*^*Cre*^; *R26R*^*DTA*^ embryos (right) and control littermates (left). Arrowheads point to sensory fiber-associated sympathetic neurons apart from the main ganglia. Scale bar = 0.5 millimeters in top panel, 0.2 millimeters in the insets. All panels show representative images of at least 3 comparisons (a pair of control and *Olig2*^*Cre*^; *R26R*^*DTA*^ embryos). Dotted boxes indicate insets. **(B)** Sagittal view of a whole mount immunostaining for TH, Neurofilament 200kD (2H3), and CD31 of E12.5 *Olig2*^*Cre*^; *R26R*^*DTA*^ embryos (first and third rows) and control littermates (second and bottom rows). Panels show respective sympathetic phenotypes at the brachial level (first two rows) and sacral level (bottom two rows). Arrowheads point at general fragmentation of the chain (middle panels), and abnormal axon growth (right panels). Scale bars = 0.5 millimeters. **(C-E)** Quantifications of volumes segmented in Imaris from wholemount stains using TH^+^ cell populations to label sympathetic cell populations in E12.5 embryos. Error bars represent standard deviation and the middle bar represents mean. **(C)** Volumes of cervical and posterior sympathetic chain ganglia in E11.5 embryo. **(D)** Quantification of the entire TH^+^ domain at E12.5. **(E)** Segmented portions of the volumes in (C), separated by anatomical regions (cervical ganglia, posterior chain, and adrenal/paraganglia). For all volumetric analysis shown in this figure, each datapoint represents the average volume of both left/right sympathetic ganglia volumes from an individual embryo. Student’s t-test was used to determine statistical significant differences between the lengths and volumes of the sympathetic ganglia of *Olig2*^*Cre*^; *R26R*^*DTA*^ versus control littermates (**p < 0.005; *p < 0.05; N.S. no significant difference detected). The test reveals significant differences in posterior chain and adrenal/paraganglia volumes between normal and motor-ablated conditions at E12.5, whether left/right ganglia are averaged per embryo (as shown) or calculated as individual datapoints (not shown).

**Figure 2:**
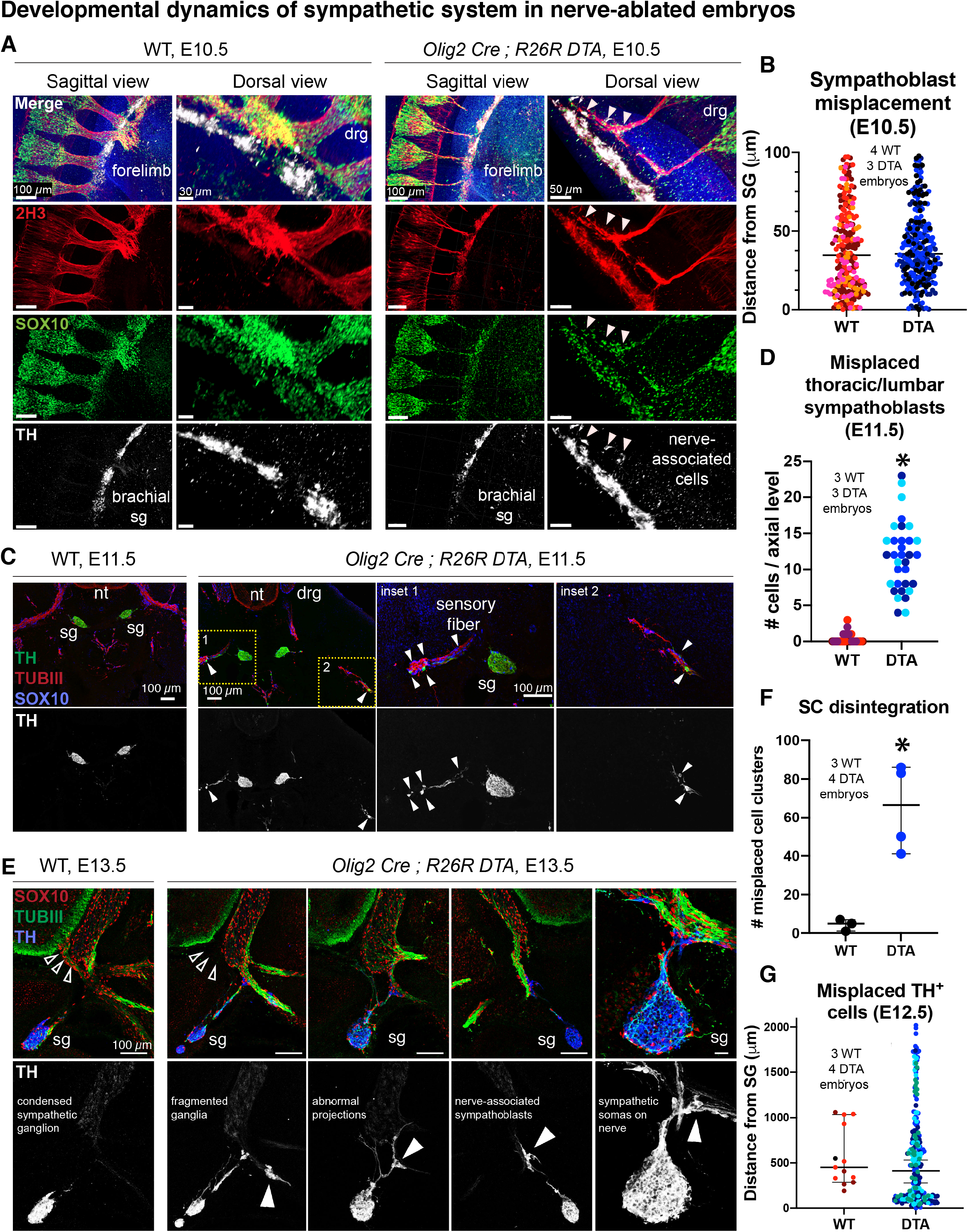
Progressive accumulation of misplaced sympathoblasts associated with sensory fibers is caused by motor ablation. **(A)** Whole mount immunostaining for peripheral nerves (2H3), SOX10, and sympathetic neurons (TH) on embryonic day 10.5 of *Olig2*^*Cre*^; *R26R*^*DTA*^ (rightmost two panels) and corresponding littermate controls (WT, leftmost two panels) at forelimb level. Arrowheads indicate sensory nerve associated Sox10+ and TH+ double positive cells. Note the thinner visceral nerve fibers following motor ablation. Sagittal view scale bars = 100 micrometers, dorsal view scale bars = 30 micrometers (left) and 50 micrometers (right). DRG: dorsal root ganglion. SG: sympathetic ganglia. **(B)** Quantification of the distance between observed sympathoblasts expressing TH and the border of the sympathetic chain (determined by automatic volume creation in Imaris). Each color of dot represents counts from a separate embryo (n=4 control and 3 mutant embryos). **(C)** Fluorescence immunostaining for SOX10, tyrosine hydroxylase (TH), and beta-3 tubulin (TUBIII) on transverse cryosections through the trunks of E11.5 *Olig2*^*Cre*^; *R26R*^*DTA*^ embryos (right panels) and control littermate (left panels). Filled arrowheads indicate TH+ sympathetic neurons associated with TUBIII+ sensory fibers. **(D)** Quantification of the number of observed misplaced sensory-nerve-attached sympathetic neurons cells per DRG at all levels between brachial and lumbar. Each color of dot represents counts from a separate embryo (n=3 control and 3 mutant embryos). **(E)** Fluorescence immunostaining for SOX10, tyrosine hydroxylase (TH), and beta-3 tubulin (TUBIII) on transverse cryosections through the trunks of E13.5 *Olig2*^*Cre*^; *R26R*^*DTA*^ embryos (right panels) and control littermate (left panels). Open arrowheads indicate the position of the ablated motor neurons in the ventral root. Filled arrowheads indicate TH+ sympathetic neurons associated with TUBIII+ sensory fibers. NT: neural tube, DRG: dorsal root ganglia, SG: sympathetic ganglia. Scale bars = 100 micrometers except rightmost panel, magnified to show soma (20 micrometers). **(F)** Quantification of the amount of TH^+^ cells observed misplaced outside the chain ganglia at E12.5. **(G)** Distribution of the distances between the observed misplaced TH^+^ cells and the borders of the chain ganglia. Each color of dot represents counts from a separate embryo (n=3 control and 4 mutant embryos).

### Motor ablation leads to misplacement of sympathoblasts aberrantly associated with sensory fibers

Although the overall morphology of the bona fide sympathetic chain was intact in E10.5 *Olig2*^*Cre*^*;R26R*^*DTA*^ embryos, SOX10^+^/TH^+^ cells were found attached to the sensory nerves in the proximity of the dorsal aorta (Figure 2A arrowheads). At this early stage when sympathoblast condensation is still in progress, the distance between individual TH^+^ cells and the sympathetic chain anlagen was not significantly different in mutant embryos compared to controls (Figure 2B). However, in E11.5 mutants, nerve associated SOX10^+^/TH^+^ cells were markedly misplaced (Figure 2C). In some cases, TH^+^ neuroblasts were attached to sensory fibers located hundreds of microns away from the dorsal aorta and sympathetic ganglia (Figure 2C, arrowheads), a large mispositioning that is unlikely to be caused by egression from the sympathetic chain. At each metameric segment between sacral/cervical ganglia in the mutants, we observed many misplaced neuronal clusters more than 50 µm away from the sympathetic chain, which is not seen in control embryos (Figure 2D). Dispersed TH^+^ cell clusters located far from sympathetic ganglia were numerous at E12.5 in mutants (Figure 2F and Figure 2G).

By E13.5, the phenotype persists in mutants, including fragmented chain ganglia, misplaced nerve-associated sympathetic neurons, and abnormal ventro-dorsal projections towards the neural tube and DRGs (Figure 2E, lower panels). The projections from misplaced cells indicate that some degree of functional maturation occurred at ectopic locations. These results were confirmed in a second genetic model of motor nerve ablation – *Hb9*^*Cre*^*;Isl2*^*DTA*^. At E14.5, mutant embryos displayed ectopic autonomic cell clusters expressing PHOX2B distributed along the sensory nerves in the vicinity of dorsal aorta (Supplementary Figure 3A). Nerve-associated TH^+^ cells were also frequently found at a distance from the sympathetic chain (Supplementary Figure 2B). Thus, in the absence of motor innervation, SOX10^+^ sympathoblast progenitors give rise to so-called “mini-ganglia” outside of the sympathetic chain that project along the branches of sensory nerves, which in wild-type embryos do not typically co-navigate extensively with sympathetic fibers between embryonic stages E10.5-E14.5. These data point to a previously unrecognized role for motor nerves in guiding the migration of sympathetic progenitor cells towards the sympathetic chain.

### Motor ablation causes misrouting of sensory nerves away from sympathetic ganglia

Because the trunk sympathetic chain is in contact with both motor and sensory nerves, the selective ablation of motor nerves might influence how autonomic progenitors, either of neural crest or SCP origin, interact with sensory axons. In line with this idea, earlier reports using *Olig2*^*Cre*^*;R26R*^*DTA*^ mice have shown that sensory nerves emanating from the dorsal root ganglia are misrouted in the absence of motor axons ^18^. First, we observed that when motor axons are depleted in E10.5 *Olig2*^*Cre*^*;R26R*^*DTA*^ embryos, a smaller fraction of viscerosensory fibers reach the sympathetic chain, while the majority either does not extend from the DRG or stay at a distance from coalescing sympathetic ganglia (Supplementary Figure 4A). Nevertheless, SOX10^+^ SCPs were found attached to the misrouted sensory nerves at multiple levels (Figure 3A). The inability of sensory nerves to properly approach the autonomic ganglia persists until E12.5 as measured by the increased distance between 2H3^+^/TH^-^ peripheral nerve fibers and the sympathetic chain in *Olig2*^*Cre*^; *R26R*^*DTA*^ mutants compared to control littermates (Supplementary Figure 4B). These peripheral nerves in motor-ablated embryos are also noticeably thinner than those in control, raising the possibility that a reduced size of migration tracts might slow migration of SOX10+ cells along the nerve (Supplementary Figure 4C). Although this experiment lacks a sensory-ablated control to tease out a specific role for sensory fibers, the current data suggests that the misplaced sympathetic neurons might result from the migration of neural crest and SCPs along improperly positioned sensory axons.

**Figure 3:**
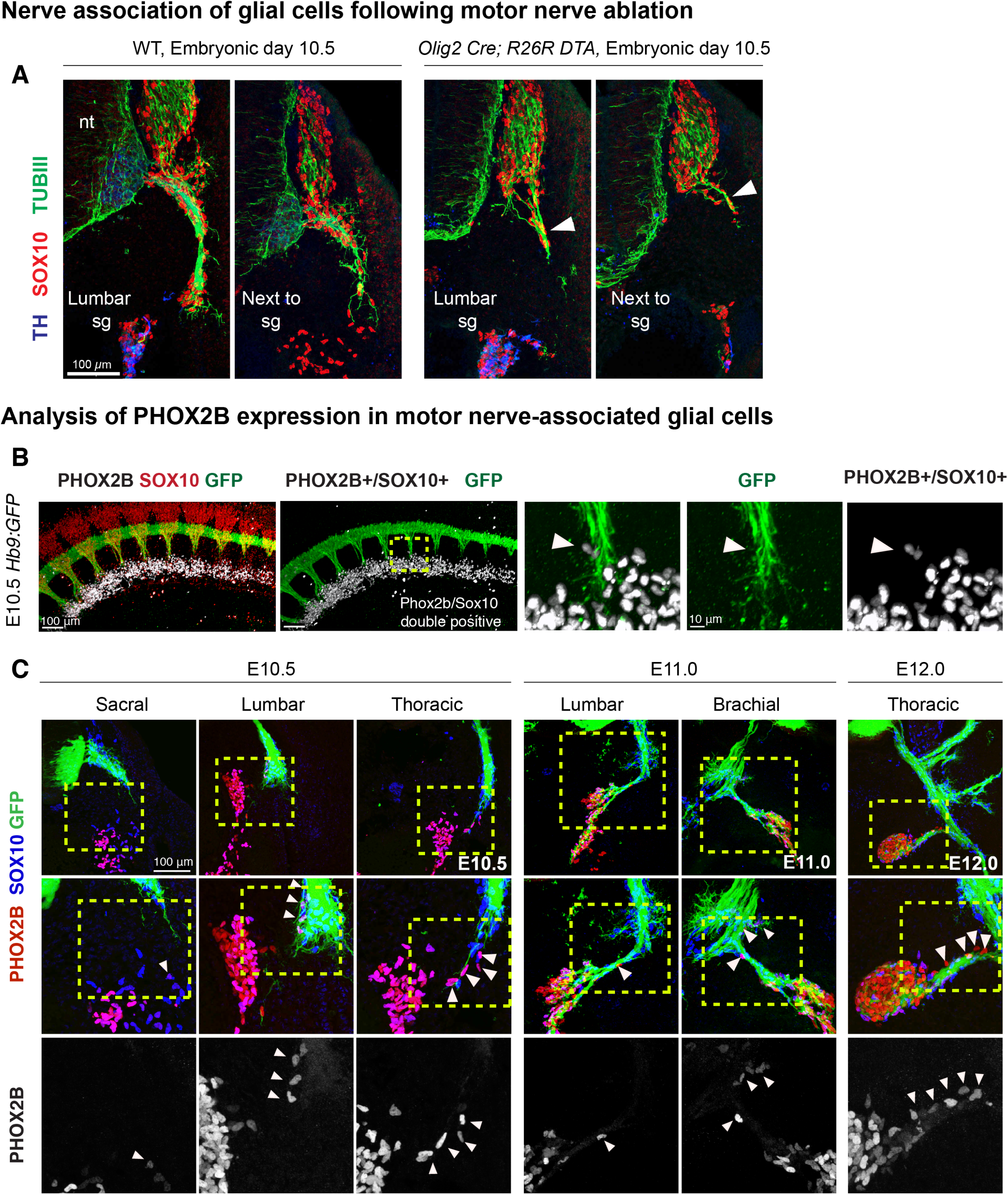
Schwann cell precursors are primed for autonomic differentiation while still migrating along motor nerves towards the sympathetic anlagen. **(A)** Immunostaining for peripheral nerves (TUBIII), SOX10, and sympathetic neurons (TH) on transverse sections through the trunk of embryonic day 10.5 of *Olig2*^*Cre*^; *R26R*^*DTA*^ (rightmost two panels) and corresponding littermate controls (WT, leftmost two panels) at the lumbar level. Arrowheads indicate sensory nerve associated SOX10+ cells. Note the reduced visceral neuron axon length following motor ablation. Scale bars = 100 micrometers. **(B**) Sagittal view of whole mount immunostaining for autonomic marker PHOX2B, glial marker SOX10, and GFP, of a stage E10.5 *Hb9:GFP* embryonic trunk. White signal is showing PHOX2B/SOX10 double-positive cells. Arrowheads in the magnified images point to a double-positive cell associated with the motor nerve. Scale bar = 100 micrometers for main panel, 10 micrometers for inset. **(C)** Transversal view of cross sections immunostained for PHOX2B, SOX10, and GFP from *Hb9:GFP* embryonic trunks. Images are ordered into a pseudo-timeseries utilizing the gradient of development along the anterior-posterior axis (lumbar, lower thoracic, upper thoracic, and brachial) across 3 time points (E10.5, E11.0, and E12.0, left to right). Insets are indicated by dotted boxes. Arrowheads indicate motor nerve associated PHOX2B+ and SOX10+ double positive cells. Scale bar = 100 micrometers.

### SCPs are primed toward sympathetic cell fate while migrating along visceral nerves

The presence of ectopic sensory nerve-associated TH^+^ cells in *Olig2*^*Cre*^*;R26R*^*DTA*^ embryos raised the question of when, and where, nerve-associated neural crest and SCP cells show early signs of sympathetic differentiation in wild-type embryos, and in particular if SCPs could trigger autonomic-specific gene expression while still on the nerve. If so, in *Olig2*^*Cre*^*;R26R*^*DTA*^ embryos, mistargeted viscerosensory fibers unable to reach the chain ganglia would disseminate autonomic-primed SCPs at incorrect locations, thus contributing to the formation of the abnormal sympathetic mini-ganglia.

First, we examined whether nerve-associated SCP express autonomic-specific genes during normal sympathetic ganglia organogenesis. At E10.5, expression of one autonomic marker, PHOX2B, in SOX10^+^ cells was largely restricted to the domain of coalescing sympathetic chain (Figure 3B). However, already at this early stage we detected expression of PHOX2B in SCP cells associated with the tips of *Hb9::GFP*^+^ visceromotor nerves approaching the sympathetic anlagen (Figure 3B, right panels, arrowheads; Figure 3C, left panels). At E11.5-E12, PHOX2B^+^ SCPs became more frequent on motor axons reaching the sympathetic chain and were observed farther away from the dorsal aorta and sympathetic anlagen as compared to the earlier stages (Figure 3C, right panels). Thus, we could detect a progressive increase in autonomic gene expression of nerve-associated neural crest and SCPs in the dorsal aorta region during normal development. Priming could be induced by potent microenvironmental signals emanating from the dorsal aorta ^14^. However, we cannot fully exclude the possibility that some sympathoblasts could migrate towards and away from sympathetic ganglia during the later stages, resulting in the growing PHOX2B expression domain along peripheral nerves.

### Ventrally migrating neural crest cells are drawn to motor axons exiting the spinal cord

Since the ectopic mini-ganglia observed in motor-less embryos were found along sensory fibers, the phenotype might be caused by misrouting of nerve-associated, neural crest-derived SCPs due to either lack of guidance signals from motor nerves or detection of inappropriate cues from the mistargeted sensory fibers. This would imply that in a normal scenario SCPs migrating along motor nerves contribute to development of the sympathetic ganglia chain, after its initial formation from the migratory neural crest. We took advantage of the antero-posterior gradient of maturation along the body axis to reconstruct developmental time series of peripheral innervation from single embryos (Supplementary Figure 5A). In E10.5 embryos, motor axons, neural crest, SCP populations, and sympathetic chain were found at distinct stages of their developmental progression at different rostro-caudal levels. At this time, SCPs were the only nerve associated SOX10^+^ cell population, whereas other SOX10^+^ cells were either freely migrating neural crest cells, or satellite glia in the forming DRG. SCPs were persistently associated with motor axons from the earliest stages of their outgrowth from the spinal cord (visible in the caudal region) until they reached the sympathetic anlagen (anterior region). Mature TH^+^ sympathoblasts were not observed prior to motor innervation (Supplementary Figure 5A). These observations led us to reason that motor nerves might guide the later waves of the neural crest migration or direct subsets of SCPs to the sympathetic ganglia. This possibility was supported by serial analysis of transverse sections through the lumbar and thoracic levels along the trunks of E10.5 *Hb9::GFP* embryos that express GFP in spinal motor neurons (Supplementary Figure 5B) ^28^. The ventral pathway of neural crest migration intersects the trajectory of motor axons exiting the neural tube and SOX10^+^ crest cells associate with motor fibers (Supplementary Figure 5B). This suggest that motor nerves “sponge” the ventral neural crest cell streams and guide a portion of the migrating neural crest/SCP towards the coalescing sympathetic chain.

### Anteroposterior gradient of PLP1-derived sympathetic chain neurons

To resolve the contribution of SCPs and satellite glial cells to the developing sympathetic ganglia, we performed lineage tracing from E10.5 to E13.5 in *PLP1*^*CreERT2*^*;R26R*^*YFP*^ embryos. This transgenic model recombines in neural crest cells before the end of axon-free migration, and in satellite glial cells and SCPs after neural crest migration is complete ^26^. At E10.5, the vast majority of the neural crest progeny is turned into neurons or nerve-associated cells along most of the trunk, with some free migrating SOX10^+^ cells evident only in the most caudal part of the embryo (Supplementary Figure 5A, right) ^26^. Since at E10.5, neural crest migration is complete in the anterior, but not posterior, regions of the embryo, *PLP1*^*CreERT2*^*;R26R*^*YFP*^-labeled cells in the cervical ganglia are supposed to derive exclusively from SCPs and satellite glia, while labeling in posterior ganglia would result from mixed PLP1^+^ populations. Consistent with this prediction, PLP1-traced cells constituted up to 30% of TH^+^ sympathoblasts in the caudal-most chain ganglia and their contribution decreased gradually to roughly 10% in the superior cervical ganglion (Supplementary Figure 5C-D). Notably, this proportion (∼ 10%) corresponds to the reduction of the sympathetic chain volume in the case of visceral motor nerve ablation (Figure 1D-E). These results support the idea that both resident PLP1^+^ progenitors and nerve-associated SCPs are sources of sympathetic neurons for the developing sympathetic chain (Supplementary Figure 5E). Of note, the relative contribution of satellite glial cells and nerve-bound SCPs cannot be resolved with cell-specific Cre drivers currently available. Overall, it seems that multiple cell sources contribute to the growing sympathetic chain after the early waves of neural crest migration ^10^, and some of them are responsible for the misplacement of sympathetic neuroblasts in the absence of motor innervation.

### Cervical ganglia malformations after motor neuron ablation

Our experiments so far suggested that cervical sympathetic ganglia and trunk sympathetic chain, which displayed distinct phenotypes in E12.5 *Olig2*^*Cre*^*;R26R*^*DTA*^ embryos (Figure 1E), might have different relationships to motor nerves during their development. Although the fragmentation phenotype was mostly evident in the posterior trunk, the shape of the superior cervical ganglion (SCG) was also affected in *Olig2*^*Cre*^; *R26R*^*DTA*^ embryos. In mutants, the cervical ganglia were grossly similar to controls at E11.5-12.5 (Figure 4A) but became gradually more deformed after E13.5 (Figure 4B-C). In wild-type embryos, the cervical sympathetic region is segmented into a large superior ganglion and a small medial ganglion. In *Olig2*^*Cre*^; *R26R*^*DTA*^ embryos, the SCG was abnormally distended (Figure 4B, upper panels) towards the level of the medial cervical ganglion (MCG). We noticed that ablation of motor neurons with *Chat*^*Cre*^*;R26*^*DTA*^ had a milder effect on cervical ganglia morphology, which became slightly elongated in a few of the mutants (Figure 4B lower panels and Supplementary Figure 6A). *Chat* expression begins in postmitotic motor neurons around E11.5 ^29,30^. We confirmed the contribution of Chat-expressing cells to motor and sympathetic neurons via lineage tracing (Supplementary Figure 6B). Thus, *Chat*^*Cre*^ becomes active days later than *Olig2*^*Cre*^, which is already functioning in motor neuron progenitors. The different severity of the phenotypes observed following DTA-mediated cell ablation might depend mostly on the timing of Cre-dependent recombination. However, one attractive hypothesis for future studies is whether early motor axon-SCG interactions, prior to the complete phenotypic maturation of motor neurons, are especially important for morphogenesis of sympathetic ganglia.

**Figure 4:**
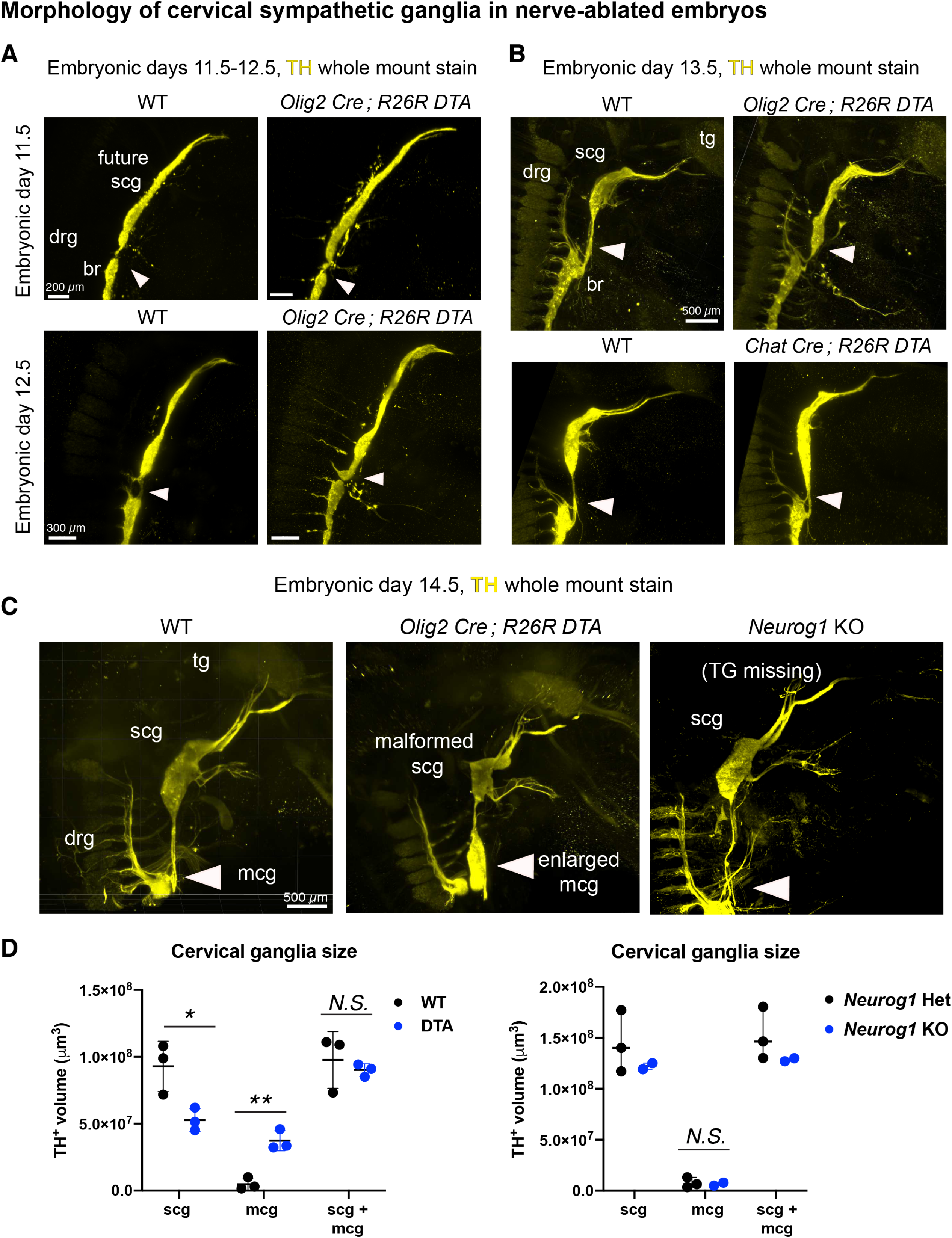
Cervical sympathetic ganglia development depends on visceromotor innervation. **(A)** Close-up sagittal view of whole mount immunostaining for TH to visualize the cervical sympathetic anlagen in E11.5 and E12.5 embryos (from the same batch of embryos shown in Figure 1C-D). Scale bars = 200 micrometers (top panels) and 300 micrometers (bottom panels) **(B)** Sagittal view of whole mount immunostaining for TH to visualize cervical sympathetic neurons in E13.5 embryos with motor ablation genotypes (right) and corresponding littermate controls (left). Motoneuron progenitor cells ablated with *Olig2*^*Cre*^; *R26R*^*DTA*^ (top) and mature motoneurons ablated with *Chat*^*Cre*^; *R26R*^*DTA*^ (bottom). WT: wildtype, tg: trigeminal ganglia, drg: dorsal root ganglia, scg: superior cervical ganglion, br: brachial level. Scale bars = 500 micrometers. **(C)** Sagittal view of whole mount immunostaining for TH to visualize cervical sympathetic neurons in E14.5 embryos with nerve ablation genotypes (right) and WT controls (left). Motoneuron progenitor cells are ablated with *Olig2*^*Cre*^; *R26R*^*DTA*^ (middle); sensory neurons are ablated in Neurog1 KO (right). WT: wildtype, tg: trigeminal ganglia, drg: dorsal root ganglia, scg: superior cervical ganglion, mcg: medial cervical ganglion. Scale bars = 500 micrometers. **(D)** Quantification of the superior cervical ganglia (scg) and medial cervical ganglia (mcg) volumes in E14.5 *Olig2*^*Cre*^; *R26R*^*DTA*^ embryos (DTA) and littermate controls (WT), as well as Neurog1 heterozygote and full KO. Each dot represents the volume of a single ganglia measured using automatically segmented surface generation in Imaris. Error bars report standard deviation, and middle bar reports mean. Student’s t-test was used to determine statistical significance (*p value < 0.05; ** p value < 0.005)

**Figure 5:**
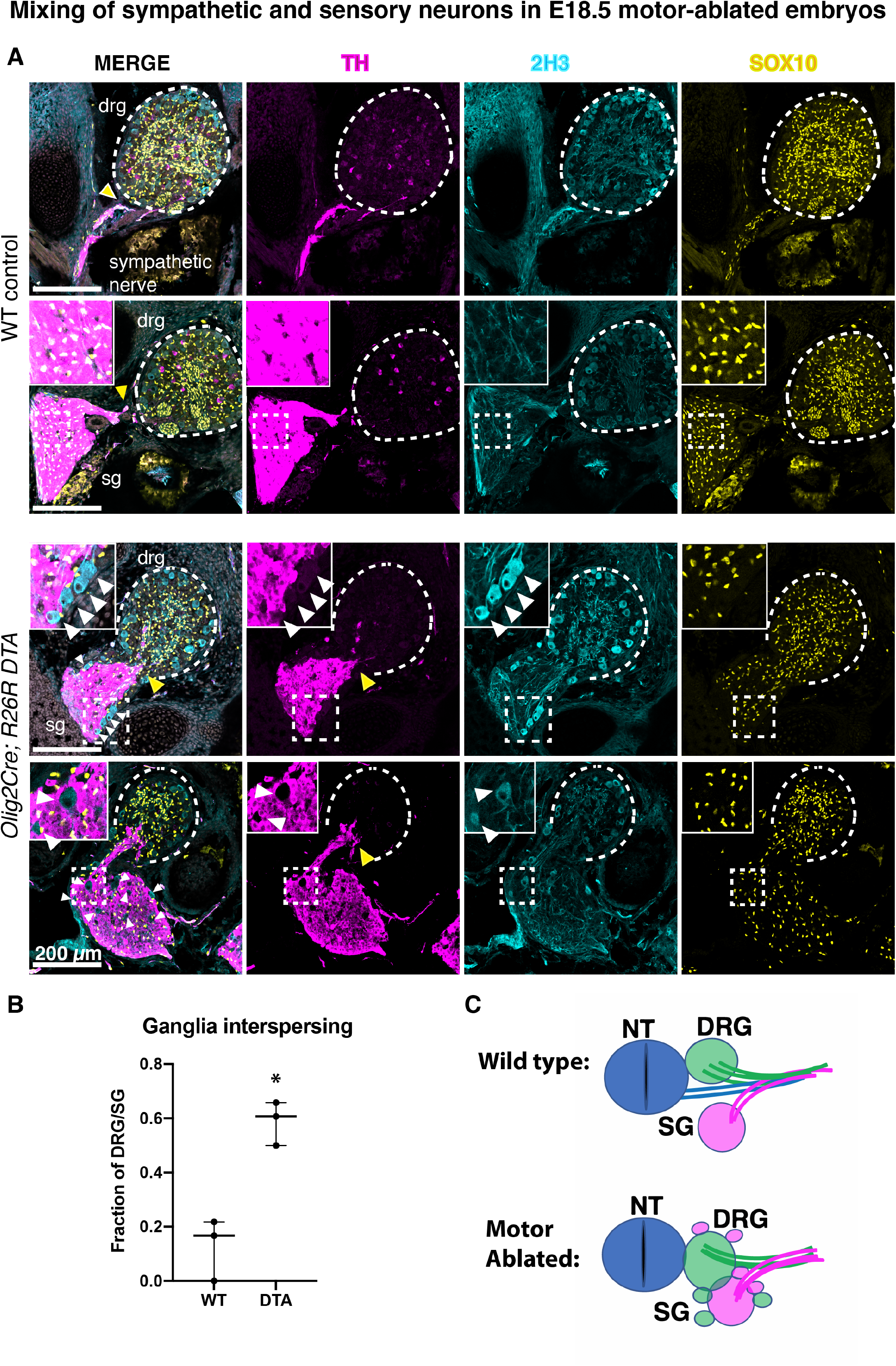
Interspersing of peripheral ganglia in late stage motor-ablated embryos. **(A)** Immunostaining for TH (magenta), SOX10 (yellow), and 2H3 (cyan) in sagittal sections of *Olig2*^*Cre*^; *R26R*^*DTA*^ embryos and wild-type littermates. The dorsal root ganglia is outlined with a dotted line. Insets emphasizing the interspersed sensory neurons in sympathetic ganglia (white arrowheads) are the regions outlined by the dotted box. Yellow arrowheads indicate the differences in peripheral ganglia proximity between wildtype and motor-ablated embryos. Scale bar = 200 micrometers. **(B)** Quantification of the fraction of images of ganglia showing aberrant organization such as misplaced sympathoblasts around the sensory ganglia, or misplaced sensory neurons in the sympathetic ganglia. Three embryos per genotype were analyzed. **(C)** Schematic illustrating the phenotype. SG=sympathetic ganglia, DRG=Dorsal root ganglia.

The misshapen ganglia phenotype progressively worsened in mutant embryos at E14.5. We measured a twofold reduction in SCG volume, which was accompanied by a corresponding increase in MCG volume (Figure 4C). Levels of cell proliferation and apoptosis did not appear to change in *Olig2*^*Cre*^; *R26R*^*DTA*^ cervical ganglia (E12.5-E13.5) compared to controls (Supplementary Figure 7), suggesting that the phenotype observed might be due to differences in collective cell movements during sympathetic ganglion development. This defect appears to depend on motor nerves but not sensory nerves, as it was not present in *Neurog1* knockout embryos in which cranial sensory innervation is disrupted ^31^ (Figure 4C, right panel).

### Motor nerves prevent abnormal intermixing of sensory and autonomic ganglia

We asked whether ganglia defects in the motor-ablated embryos would persist at later time points. Since *Olig2*^*Cre*^; *R26R*^*DTA*^ pups die at birth ^18^, we analyzed ganglia morphology at E18.5. Unexpectedly, we found that in mutant embryos sympathoblasts expressing high levels of TH (in contrast to typical sparse, low-TH sensory neurons in normal DRG) were inappropriately located around and within the dorsal root ganglia and, conversely, that sensory neurons were misplaced in the sympathetic ganglia (Figure 5A) to the degree that sensory and autonomic ganglia appeared to be mixed and merged at brachial levels. This phenotype was highly penetrant, as it was found in all DTA embryos analyzed and never in controls (Figure 5B; n=3 embryos assessed per genotype, 12-38 sections assessed per embryo). Thus, motor nerves appear to be important for partitioning and preserving the integrity of sensory and sympathetic ganglia (schematic summarizing result in Figure 5C).

The disruption of sympathetic development caused by motor nerve ablation suggests that signals from preganglionic nerves may instruct neural crest cell development. To examine this possibility, we re-analyzed publicly available single-cell transcriptomes of neural crest cells ^32^ and spinal motor neurons ^33^ from embryonic stages E9.5-E10.5 (Figure 6A) and used CellChat algorithm to predict ligand-receptor interactions between these cell types ^34^. Different subtypes of motor neurons, including preganglionic neurons (PGC) expressed ligands that were predicted to signal to neural crest cells expressing cognate receptors (Figures 6B and 6C). This analysis identified five candidate pathways predicted to mediate motor neuron-to-neural crest cell signaling: Class 3 Semaphorin signaling (SEMA3, Figure 6D), Platelet derived growth factor (PDGF, Figure 6E), Energy Homeostasis-associated (ENHO, Figure 6F), Neuregulin (NRG1, Figure 6G), and Growth-arrest specific (GAS6, Figure 6H). While the amounts of predicted crosstalk between motor neuron clusters varied among the candidate pathways, in all cases the putative signals were expressed by motor neurons and the receptor was also expressed by neural crest cells (Figure 6I).

**Figure 6:**
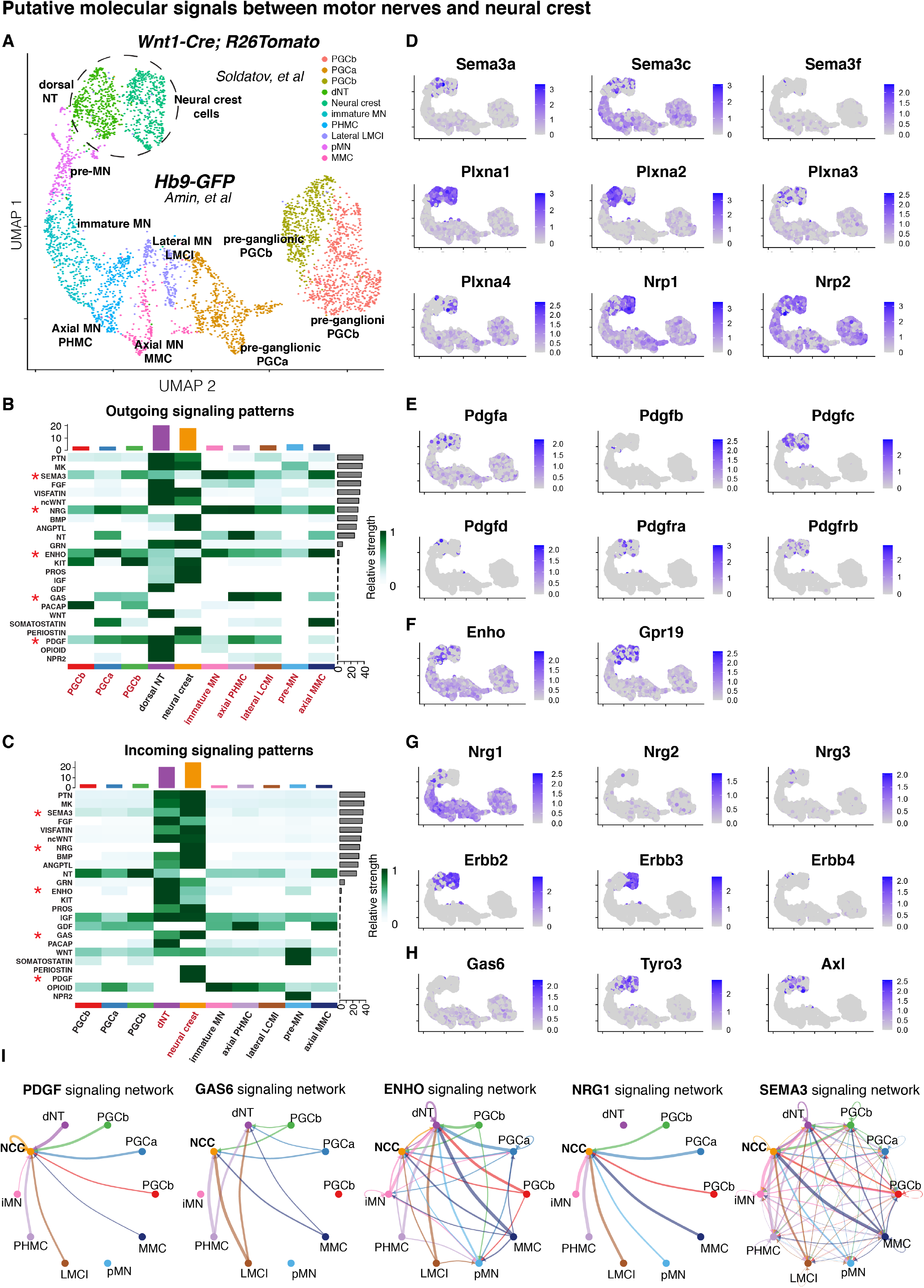
Putative signals between motoneurons and neural crest. **(A)** Joint UMAP embedding of publicly available and previously published single-cell transcriptomics datasets from E9.5 trunk neural crest cells (traced via Wnt1-Cre; tdTomato) and E10.5 spinal motor neurons (labeled via Hb9-GFP). Different colors of dots represent the previously annotated cell type clusters. The dotted circle indicates the position of cells derived from the Wnt1 tracing (Soldatov et al), and the remaining cells are derived from the Hb9-GFP labeling. Abbreviated motor neuron cell type names (MMC, LMCI, PGCa, PGCb, PHMC) refer to the originally published cluster names (Amin et al). MN = Motor neuron. NT = neural tube. **(B)** Heat map of predicted outgoing signals from each cluster in the integrated dataset. Dark green indicates expression of signaling ligands from a particular pathway. Pathways, identified on the left side, are sorted by relative strength of the gene expression signature (shown on the right side). Motoneuron clusters are highlighted in red. **(C)** Heat map of predicted incoming signals from each cluster in the integrated dataset. Dark green indicates expression of signaling receptors from a particular pathway. Pathways, identified on the left side, are sorted by relative strength of the gene expression signature (shown on the right side). The neural crest cluster is highlighted in red. **(D-H)** Expression maps of signals and cognate ligands that pertain to the top 5 candidate signaling pathways predicted by CellChatDB. Expression profiles are taken from the joint UMAP found in (A). The darker purple regions indicate increased gene expression relative to other cells in the dataset. **(D)** Expression maps of various Semaphorin pathway members. **(E)** Expression maps of various Platelet Derived Growth Factor pathway members. **(F)** Expression maps of various Energy Homeostasis-related pathway members. **(G)** Expression maps of various Neuregulin pathway members. **(H)** Expression maps of various Growth Arrest Specific 6 pathway members. **(I)** Chord plots showing the predicted directionality of signaling between each cluster, for the top 5 candidate signaling pathways. Arrows point towards the signal receiving cells, and away from the signal-producing cells.

Because the Semaphorin pathway, the top candidate from our ligand-receptor analysis, has been linked to sympathoadrenal and neural crest development by previous studies ^35 36^, we analyzed the role of Class 3 Semaphorin signaling in the formation of sympathetic chain ganglia. *Sema3A, Sema3F* and *Sema3C* were detected in the thoracic motor columns at E11.5 by *in situ* hybridization (Figures 7A and Supplementary Figure 8A), also in agreement with previous knowledge ^37^. Neural crest-specific deletion of the Semaphorin receptor Neuropilin 1 results in sympathetic chain defects at E12.5, similar to the *Olig2*^*Cre*^; *R26R*^*DTA*^ phenotype with misplaced TH^+^ neurons near to the forming chain (Figure 7B and 7C). To check if SEMA3C (highest expression in motoneurons based on the single cell data: Figure 7E and ^38^) signals from the motor nerves to sympathetic progenitors, we checked the sympathetic phenotype in the full knockout of *Sema3C* and found that TH^+^ cells and their positioning were not affected (Supplementary Figure 8B). Therefore, we concluded that SEMA3C does not play a major role in sympathetic neuron positioning. At the same time, we found a mild sympathetic phenotype in *Nrp1*^*SEMA3*^ which are special mutants of Neuropilin1 deficient in binding Semaphorin ligands. Similarly, we found the mild disruption of a sympathetic chain in NRP2 knockout mutants, (Supplementary Figure 8C), suggesting that other Neuropilin-Semaphorin interactions might be involved. To make sure that Semaphorin and not VEGF signaling is involved in developing sympathetic phenotype, we took advantage of NRP1^SEMA3^ NRP2^KO^ double mutant embryos, where all class-3 Semaphorin signaling is abolished, while VEGF signaling is still active ^39,40^. In those embryos, the defects in sympathetic ganglia were severe (Figure 7D-I). In line with this, we observed no deficits in sympathetic ganglia development in Olig2Cre-VEGF KO embryos (data not shown).

**Figure 7:**
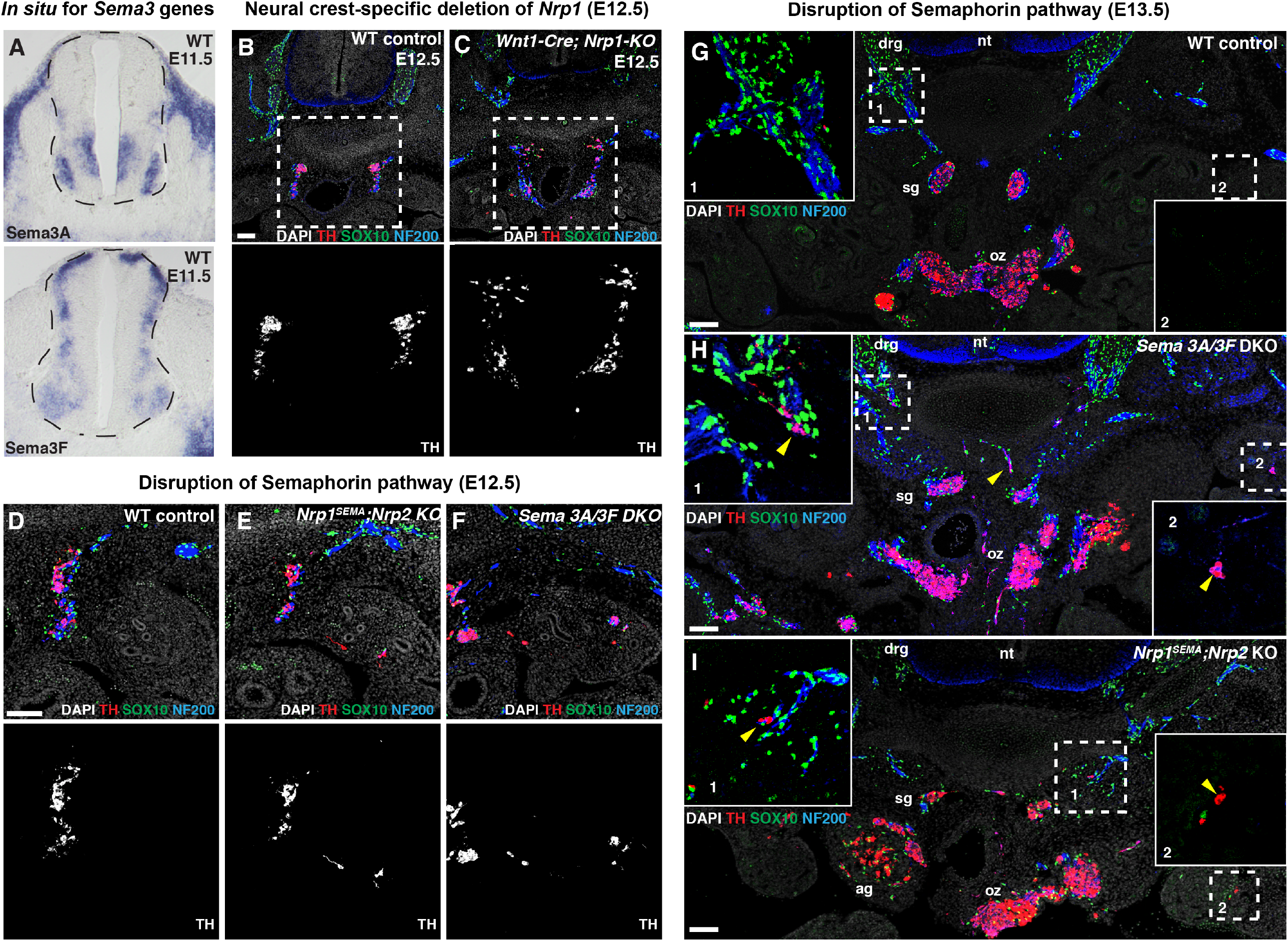
Sema3 signaling via Nrp1 regulates sympathoblast positioning. **(A)** *In situ* hybridization of cryosections of mouse embryonic trunk neural tube at E11.5. Top panel, expression of Semaphorin 3A. Bottom, expression of Semaphorin 3F. **(B-C)** Top panels, immunofluorescence staining of TH (red), Sox10 (green), and NF200 (blue) as well as DAPI (grey) to mark sympathetic neurons, glial, peripheral nerves and nuclei, respectively in (B) a control littermate or (C) a *Wnt1*^*Cre*^; *Nrp1*^*flox/flox*^ mouse embryo. TH channel is isolated in bottom panels. **(D-F)** Sympathetic ganglia in a (D) wildtype control embryo, (E) *Nrp1*^*SEMA/SEMA*^; *Nrp2*^*-/-*^ mutant embryo, and (F) Sema3A/Sema3F double knockout mutant embryo at embryonic day 12.5. Top panels, immunofluorescence staining of TH (red), Sox10 (green), and NF200 (blue) as well as DAPI (grey) to mark sympathetic neurons, glial, peripheral nerves and nuclei, respectively. TH channel is isolated in bottom panels. **(G-I)** Sympathetic ganglia in a (G) wildtype control embryo, (H) Sema3A/Sema3F double knockout mutant embryo and (I) *Nrp1*^*SEMA/SEMA*^; *Nrp2*^*-/-*^ mutant embryo, at embryonic day 13.5. Immunofluorescence staining of TH (red), Sox10 (green), and NF200 (blue) as well as DAPI (grey) to mark sympathetic neurons, glial, peripheral nerves and nuclei, respectively. Insets are shown without DAPI. Yellow arrowheads mark a few examples of misplaced sympathoblasts. Scale bars = 20 micrometers.

The above experiments return us to the role of *Sema3A* and *Sema3F –* the remaining Semaphorins, which are expressed in the motoneurons. The double knockout of *Sema3A* and *Sema3F* revealed a severe phenotype, showing numerous and widespread misplaced sympathetic neurons at E12.5 (Figure 7D-7F) and E13.5 (Figure 7G-7I), which was highly similar to the phenotype observe in motor nerve ablated *Olig2*^*Cre*^; *R26R*^*DTA*^ embryos. These results are consistent with a model in which Semaphorin signals from motor neurons direct a subset of neural crest cells to migrate along axons towards the sympathetic chain ^35,36^.

Overall, our mouse genetic studies demonstrate that motor nerves influence the composition, structure, and projections of sympathetic ganglia. The results obtained from genetic ablation of motor nerves, spatial analysis of primed progenitor populations, lineage tracing and 3D reconstruction experiments, indicate that motor nerves assist the development of the sympathetic ganglia by affecting the allocation of motor nerve-associated progenitor cells, possibly via Semaphorin signaling. Given the fact that nerve-associated SCPs represent a minor progenitor source for sympathetic neuron formation ^10 25^, it might be that in the absence of a migratory template provided by motor nerves, the PHOX2B+ SCPs (Figure 3C) generate misplaced clusters of sympathoblasts. In turn, this results in fragmented sympathetic chain, and ectopic sympathetic mini-ganglia associated with the viscerosensory branches near the dorsal aorta. Finally, visceral motor nerves are instructive for the development of sympathetic innervation patterns. In the absence of the preganglionic motor innervation, sympathetic neurons project erroneously towards the dorsal neural tube and DRGs, ultimately causing the fusion of these ganglia and abnormal intermixing of autonomic and sensory neurons.

## DISCUSSION

Besides transmitting information through synaptic contacts, peripheral axons influence the innervated tissues in a variety of ways, by secreting morphogenic navigating signals ^41^, organizing nerve tracts through axon-axon interactions ^42 43^, guiding and remodeling blood vessels ^44 45 38^ and delivering neural cells and glia cell precursors to their final destination during embryonic development ^26 7^. Because the PNS is multifunctional, the formation of innervation patterns has far-reaching consequences for organismal physiology ^46^. During organogenesis, peripheral nerves navigate throughout the embryo in response to multiple guiding cues from surrounding developing tissues and establish intertwined and co-dependent innervation patterns along the anteroposterior axis ^47^. The co-dependency of different nerve types reflects a parsimonious solution for neural wiring, and represents an appealing developmental mechanism for generating evolutionary novelty. Here, we addressed whether assembly of the sympathetic autonomic system depends on a pre-existing template of nerve fibers laid down by pioneering visceral motor neurons of the spinal cord. We report that the development of the sympathetic ganglia system requires an intact pattern of motor innervation. When motor nerves – but not sensory fibers – are selectively ablated in genetic mouse models, the sympathetic ganglia chain is disorganized and a number of sympathoblasts appear at incorrect locations along sensory nerves, suggesting that sympathetic progenitor cells migrating along peripheral motor tracts have been misplaced.

The rearrangement may result from missing signals normally provided by motor axons. The phenotype observed in the absence of motor axons was remarkably reminiscent of the defects that arise in sympathetic chain when Semaphorin/Nrp1 signaling is impaired in mutant embryos. Sema3/NRP pathway is required for sympathetic nervous system development ^35^ and for placing of chromaffin cell precursors in the adrenal medulla following visceral motor nerves ^36^. Motor neurons express multiple Semaphorins ^37^ and use them to regulate guidance receptor activity in an autocrine fashion ^48^, and as paracrine signals to control the interactions between developing motor axons and the cells in the innervated tissues, including vascular endothelial cells ^38^. Our results support the possibility that Sema3A and Sema3F ligands released by extending preganglionic motor nerves might orchestrate the migratory patterns, spatial organization and differentiation of sympathoblasts. However, since activation of Sema3/Neuropilin signaling within motor neurons has been implicated in peripheral nerve targeting to muscles and adrenal primordia ^49 50 36^, a contribution of motor axonal misrouting to sympathoblast disorganization cannot be excluded. Given that our Sema3/Neuropilin mutants showed misplaced sympathoblasts both on and off the nerve, the role of this pathway as a guidance mechanism for nerve-associated cell migration represents an interesting and complex subject of future investigation.

We found that SCPs become primed to undertake an autonomic fate while still migrating along visceral motor nerves toward the sympathetic anlagen. In our experimental paradigms, the ablation of motor nerves depletes the local pools of nerve-associated or nerve-influenced progenitors and results in misplacement of the remaining niches onto the spared and misrouted sensory nerve fibers. This change in nerve-type substrate leads to size reduction of the sympathetic chain and formation of ectopic sympathetic ganglia away from the traditional locations in the proximity of the dorsal aorta. The displacement of sympathetic ganglia is further brought about by disruption of sensory axon pathfinding in the absence of the motor nerve template. Motor neurons are the first to establish nerve tracts in the PNS and their trajectory are followed by sensory and postganglionic autonomic fibers ^18,43^. We find that as soon as the pioneering axons of motor neurons exit the spinal cord, they “sponge” the stream of freely migrating neural crest cells, which become attached to the motor fibers. Thus, early-extending motor axons function as an adhesive barrier that intersects the ventral migratory path of neural crest cells. The time of initial motor axon outgrowth from the spinal cord at each rostrocaudal segment coincides with a switch from free-to axon-associated migration of neural crest derivatives. Along the body axis, the temporal overlap between the initial establishment of motor system connectivity and neural crest migration determines the amount the nerve-associated progenitors allocated to different organs and, consequently, defines the relative contribution of migratory neural crest-independent and SCP-dependent modes of PNS morphogenesis. Even minor perturbations in the timing of motor nerve outgrowth or neural crest migration may significantly affect the composition of precursor types supplied to each organ.

We noticed that axonal navigation of the emerging sympathetic neurons was impaired in the absence of visceral motor nerves. In this experimental setting, both properly positioned and ectopic sympathetic neurons displayed enhanced axonal growth and mis-projected towards the ventral roots and dorsal root ganglia, away from their canonical path. Ultimately, at later developmental stage, the sympathetic axons project dorsally into DRGs, and sympathetic and sensory ganglia start to fuse causing inappropriate mixing of different neuronal types. These phenotypes point to a previously unappreciated inhibitory effect of preganglionic motor fibers on the early projection of sympathetic nerves. The identification of motor-derived signals that influence sympathetic axon guidance is complicated by the fact that this effect could be in part mediated by sensory axons, which also depend on pioneering motor axons to establish their connectivity patterns ^18^. Unmasking the molecular mechanisms underlying repulsive interactions between visceromotor and sympathetic fibers represents an important biological question for future studies.

Overall, our results suggest that motor nerves provide repulsive navigational cues to sympathetic innervation and also provide a niche or migratory substrate for a portion of progenitors contributing to the development of the sympathetic chain. Future studies will clarify the exact nature of repulsive molecular signals provided by motor fibers and the molecular phenotypes of progenitors interacting with motor nerves during sympathetic system development.

## METHODS

### Mouse lines

All animal work was permitted by the Ethical Committee on Animal Experiments (Stockholm North committee) and conducted according to The Swedish Animal Agency’s Provisions and Guidelines for Animal Experimentation recommendations. *R26R*^*Tomato*^ mice were ordered from The Jackson Laboratory (stock number 007914). *Plp1*^*CreERT2*^ mice were received from U. Suter laboratory (ETH Zurich, Switzerland) (http://www.informatics.jax.org/allele/MGI:2663093). *R26R*^*YFP*^ mice were received from The Jackson Laboratory (stock number 006148, full strain name B6.129X1-Gt(ROSA)26Sortm1(EYFP)Cos/J). *HB9*^*Cre*^ (also known as Mnx1Cre) mice were received from The Jackson Laboratory, stock number 006600 (full strain name B6.129S1-Mnx1tm4(cre)Tmj/J). *Isl2*^*DTA*^ mice were received from The Jackson Laboratory, stock number 007942 (full strain name B6.Cg-Isl2tm1Arbr/J). *R26R*^*DTA*^ mice were received from The Jackson Laboratory, stock number 006331 (full strain name Gt(ROSA)26Sortm1(DTA)Jpmb/J). *Chat*^*Cre*^ mice were received from K. Meletis lab (Karolinska Institutet) also available from the Jackson Laboratory, stock number 006410 (full strain name B6;129S6-Chattm2(cre)Lowl/J). All tracing experiments using *Plp1*^*CreERT2*^ ; *R26R*^*YFP*^ were performed using heterozygotes for both the Cre and the reporter *R26R*^*YFP*^ or *R26R*^*Tomato*^. Mouse mutants deficient in Semaphorin and Neuropilin signaling (*Wnt1-Cre Nrp1*^*fl/fl*^, *Sema3A*^*-/-*^, *Sema3C*^*-/-*^ *Sema3F*^*-/-*^, *Nrp1*^*SEMA/SEMA*^ *Nrp2*^*-/-*^) have been described previously ^35,36,38,39^. For all experiments, the day the plug was detected was considered E0.5. Tamoxifen (Sigma, T5648) was dissolved in corn oil (Sigma, 8267) and delivered via intra peritoneal (i.p.) injection to pregnant females and pups (0.05 mg/g body weight).

### Motoneuron ablation experiments

For targeted ablation of the pre-ganglionic neurons, *Isl2*^*DTA*^ or *R26R*^*DTA*^ mice were bred to *Hb9*^*Cre*^, mice, *Chat*^*Cre*^ mice, or *Olig2*^*Cre*^ mice to generate experimental *Hb9*^*Cre*^*/+;Isl2*^*DTA*^*/+* and control *Isl2*^*DTA*^/+ embryos, *Chat*^*Cre*^/+;*R26R*^*DTA*^/+ and control *R26R*^*DTA*^/+ embryos, or *Olig2*^*Cre*^/+;*R26R*^*DTA*^/+ and control *R26R*^*DTA*^/+ embryos. Sample size was determined by availability of mutant embryos from each litter. Images shown in the figures represent comparisons between at least three mutant and three control embryos, for each developmental stage from E10.5-E14.5.

### Immunohistochemistry

Embryos were harvested and fixed overnight using 4% paraformaldehyde dissolved in a PBS buffer (pH 7.4) at 4°C. Samples were washed in PBS at 4°C for one hour and cryopreserved by submerging at 4°C for 24 hours in 30% sucrose, diluted in PBS. The samples were then embedded using OCT media, frozen on dry ice, and stored at -20°C. Tissue blocks were sectioned on an NX70 cryostat at a section thickness of 14-40 μm. Slides were stored at -20°C after drying at RT for one hour. For antigen retrieval, slides were immersed in 1x Target Retrieval Solution (Dako, S1699) for 1 hour, pre-heated to 90°C. Sections were washed three times in PBS containing 0.1% Tween-20 (PBST), incubated at 4°C overnight with primary antibodies diluted in PBST in a humidified chamber. Finally, sections were washed in PBST and incubated with secondary antibodies and Hoescht stain diluted in PBST at RT for two hours, washed again three times in PBST and mounted using Mowiol mounting medium (Dako, #S3023). Rabbit anti-TH (1:1000, Pel-Freez Biologicals, #P40101-150), sheep anti–TH (1:2000, Novus Biologicals, #NB300-110), chicken anti-TH (1:500, Abcam, #ab76443), mouse monoclonal antibIII tubulin (1:500, Promega, #G712A), mouse anti-tubulin bIII (1:1000, Abcam, #ab7751), goat anti-GFP (1:500, Abcam, #ab6662), chicken anti-GFP (1:500, Aves Labs Inc., #GFP-1020), rabbit anti-GFP (1:5000, Thermo/LifeTech, #A6455), rabbit anti-DsRed (1:300, Clontech, #632496), chicken anti-mCherry (1:1000, EnCor Biotech, #CPCA-mCherry), goat-anti-PHOX2B (R&D, 1:1000, #AF4940), mouse anti-Neurofilaments (1:200, clone 2H3, DSHB), goat anti-SOX10 (1:500, Santa-Cruz, #sc-17342), rabbit anti-Sox10 (1:2000, Abcam, # ab155279), rabbit anti-PRPH (1:500, Chemicon, #AB1530), rabbit anti-KI67 (1:500, Thermo Scientific, #RM-9106), rat anti-PECAM (1:300, BD Pharmingen, #553370). DAPI (Thermo Fisher Scientific, 1:10,000, #D1306) was used concomitantly with secondary antibodies diluted in PBST buffer. For detection of the primary antibodies, secondary antibodies raised in donkey and conjugated with Alexa-488, -555 and -647 fluorophores were used (1:1000, Molecular Probes, Thermo Fisher Scientific).

### *In situ* hybridization

For HCR in Figure 7, probes for Sema3A and Sema3C were ordered directly from Molecular Instruments and the procedure was performed according to the protocol described online (https://files.molecularinstruments.com/MI-Protocol-RNAFISH-FrozenTissue-Rev3.pdf) using tissue cryosections from the trunks of day 11.5 mouse embryo. Briefly, slides were pre-treated via a fixation in paraformaldehyde at 4°C, dehydration with an ethanol gradient, and a 10 ug/uL proteinase K digestion for 10 minutes before transcript detection. Then, slides were hybridized to 0.4 pmol probes diluted in 100uL probe hybridization buffer (Molecular Instruments) at 37 °C overnight. After unbound probe was washed away with probe wash buffer (Molecular Instruments) the bound probes were amplified using snap-cooled H1 and H2 hairpins overnight in a dark humidified chamber. After washing excess amplification solution away using SSCT, slides were mounted and imaged on a Zeiss LSM980-Airyscan confocal microscope. For traditional *in situ* hybridization in Supplementary Figure 6, embryos were fixed overnight in cold 4% paraformaldehyde/PBS, washed in PBS, dehydrated in methanol and stored at − 20 °C. Then, *in situ* hybridization was performed using digoxigenin-labelled riboprobes transcribed from plasmids containing Sema3a and Sema3f cDNAs.

### Whole mount immunostaining of mouse embryos

After incubation of embryos with secondary antibodies and washing steps, the embryos were optically cleared using two different methods according to their size. Embryos at or younger than embryonic day E11.5 were cleared in BABB solution (1 part benzyl alcohol / 2 parts benzyl benzoate) for one hour with rotation before imaging on a confocal LSM800 microscope (Adameyko *et al*., 2009). Embryos at or older than embryonic day E12.5 were imaged using the CUBIC method (3-7 days in CUBIC1 at 37 degrees, 2-3 days in CUBIC2 solution at room temperature) before imaging on a Zeiss light sheet microscope (Susaki et al., 2015). 3D reconstructions of the sympathetic ganglia were performed using IMARIS software based on segmentations of the TH/PHOX2B staining. In some cases, autofluorescence was used to aid the identification of the vasculature, the outline of the embryo, and the neural tube.

### Microscopy

Images were acquired using LSM800 Zeiss confocal microscope or Zeiss Z1 light sheet microscope. Confocal microscope was equipped with 10x/0.45, 20x/0.8, 40x/1.2 and 63x objectives. Laser lines used for excitation included 405nm, 488nm, 561nm, and 640nm. All images using the light sheet were taken with 5X/0.16 air objective, using 405nm, 488nm, 561nm, and 638 nm for excitation. Light sheet images were acquired in the .czi format and processed by stitching with Arivis Vision 4D, down-sampling (1:2 in the XY plane) in ImageJ, conversion to .ims files using Bitplane Imaris File Converter, and downstream analysis with Bitplane IMARIS (9.5). Confocal images were acquired in .czi format and analyzed in FIJI.

### Image analysis for ganglion volume quantification

Volumetric quantification of pelvic ganglia was performed using Bitplane Imaris 9.5 with the surface generation tool and a fixed intensity threshold for all samples. Volumetric quantification of sympathetic chain volumes was performed using Bitplane Imaris 9.5 using the surface generating tool. 5 to 7.5 µm smoothing was used for all light sheet images, 1.25 µm smoothing was used for confocal images. Manual segmentation was used to separate cervical ganglia, sympathetic chain, adrenals, and misplaced sympathetic neurons before surface generation. Signal intensity thresholds, which were automatically recommended by the Imaris software for each whole unsegmented image, were used for surface generation. A two-sided Student’s t-test was performed to determine statistically significant differences between the volumes of the sympathetic chains. At least three embryos were used for the sympathetic ganglia volume analysis, per stage, for all comparisons between motor-ablated and control embryos.

### Image analysis of cell misplacement

From the three-dimensional fluorescence micrographs of E11.5 and E12.5 motor-ablated and control embryos stained for TH and 2H3, surfaces including only condensed sympathetic chain ganglia were generated by manual segmentation. Manual segmentation was also used to create surfaces including misplaced sympathoblasts, to avoid counting enteric neurons or adrenal cells. For E10.5 images, the surface representing the sympathetic chain was generated automatically, and enteric cells were excluded by filtering out TH-positive cells farther than 100 µm from the sympathetic anlagen. Spot detection algorithm in Imaris was used for cell counting (expected object XY size = 10 µm, Z correction = 20 µm).

### Image analysis of lineage tracing

Quantification of lineage tracing experiments was performed using FIJI. Multicolor, multi-tile, Z-stack images in CZI format with channels for DAPI, TH, and YFP were converted into 8-bit, and all Z-slices were combined using a maximum intensity projection. Binarization of images was based on manual thresholding per image, after it was determined that a fixed threshold per embryo gave similarly trending results across the anteroposterior axis (data not shown). Image calculation in FIJI was used to find single, double, and triple positive regions. The ratio between the amount of triple positive YFP^+^ TH^+^ DAPI^+^ objects and the amount of double positive TH^+^ DAPI^+^ objects was used to determine the percent of traced neurons in the sympathetic ganglia. Over 1000 cells were counted per axial region per embryo, and the graphical results are from the averages of four analyzed embryos from two different litters.

### Cell-cell interaction analysis

Single cell transcriptomics datasets used for the study are available for download from (http://pklab.med.harvard.edu/ruslan/neural.crest.html) and ArrayExpress accession: E-MTAB-10571. Standard pre-processing workflow such as QC, cell selection, data normalization, identification of highly variable features, scaling, dimensional reduction, clustering, and integration of the datasets was performed using Seurat according to the instructions provided (https://satijalab.org). Cell interaction analysis was performed using the R package CellChat. A subset of CellChatDB (“Secreted signaling”) was used to seek cell-cell interactions between cell clusters in the integrated motor-crest Seurat object (https://github.com/sqjin/CellChat).

### Data, code, materials, and protocol

Information provided in the text, figures and supplementary information contained in the present manuscript is sufficient to assess whether the claims of this study are supported by the evidence. Datasets, code, materials, and protocols used for this research project are available to interested parties upon request.

## ACKNOWLEDGEMENTS

AGE was supported by StratNeuro SRP Postdoctoral Research Fellowship and F32 NRSA NIH/NIDCR. IA was supported by ERC Consolidator grant STEMMING-FROM-NERVE and ERC Synergy Grant KILL-OR-DIFFERENTIATE, Swedish Research Council, Knut and Alice Wallenberg Foundation, Bertil Hallsten Research Foundation, Cancerfonden, Paradifference Foundation. DB is supported by European Research Council Starting Grant 335590 and a Career Development Award from the Giovanni Armenise-Harvard Foundation.

## COMPETING INTERESTS STATEMENT

The authors declare no competing interests.

## Notes

### Competing Interest Statement

The authors have declared no competing interest.

