## Supplement for "Motor nerves direct the development of the sympathetic nervous system"

Motoneuron ablation efficiency of *Olig2Cre; R26R DTA* embryos

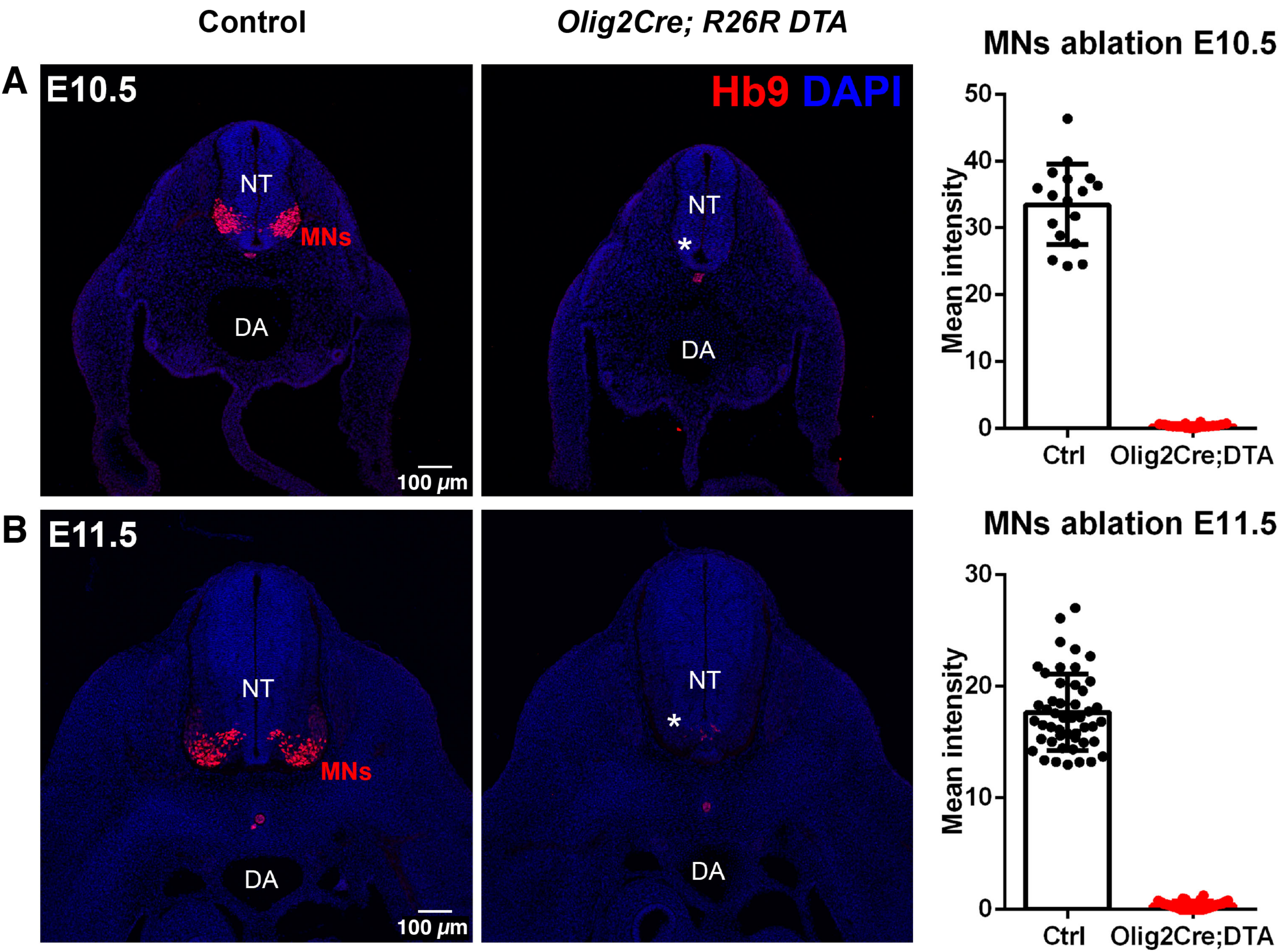


Supplementary Figure 1

**Supplementary figure 1: Olig2-DTA is an efficient and specific mouse model for motor nerve ablation. (A-B)** Immunostaining for HB9 (red) in *Olig2$^{Cre}$; R26R$^{DTA}$* embryos and wild-type littermates. Embryos were checked at stage E10.5 (A) and E11.5 (B). NT: neural tube, DA: dorsal aorta. Scale bar = 100 micrometers. Right panels show quantifications for the loss of HB9$^{+}$ cells.

Sympathoadrenal development in *Olig2Cre; R26R DTA* embryos

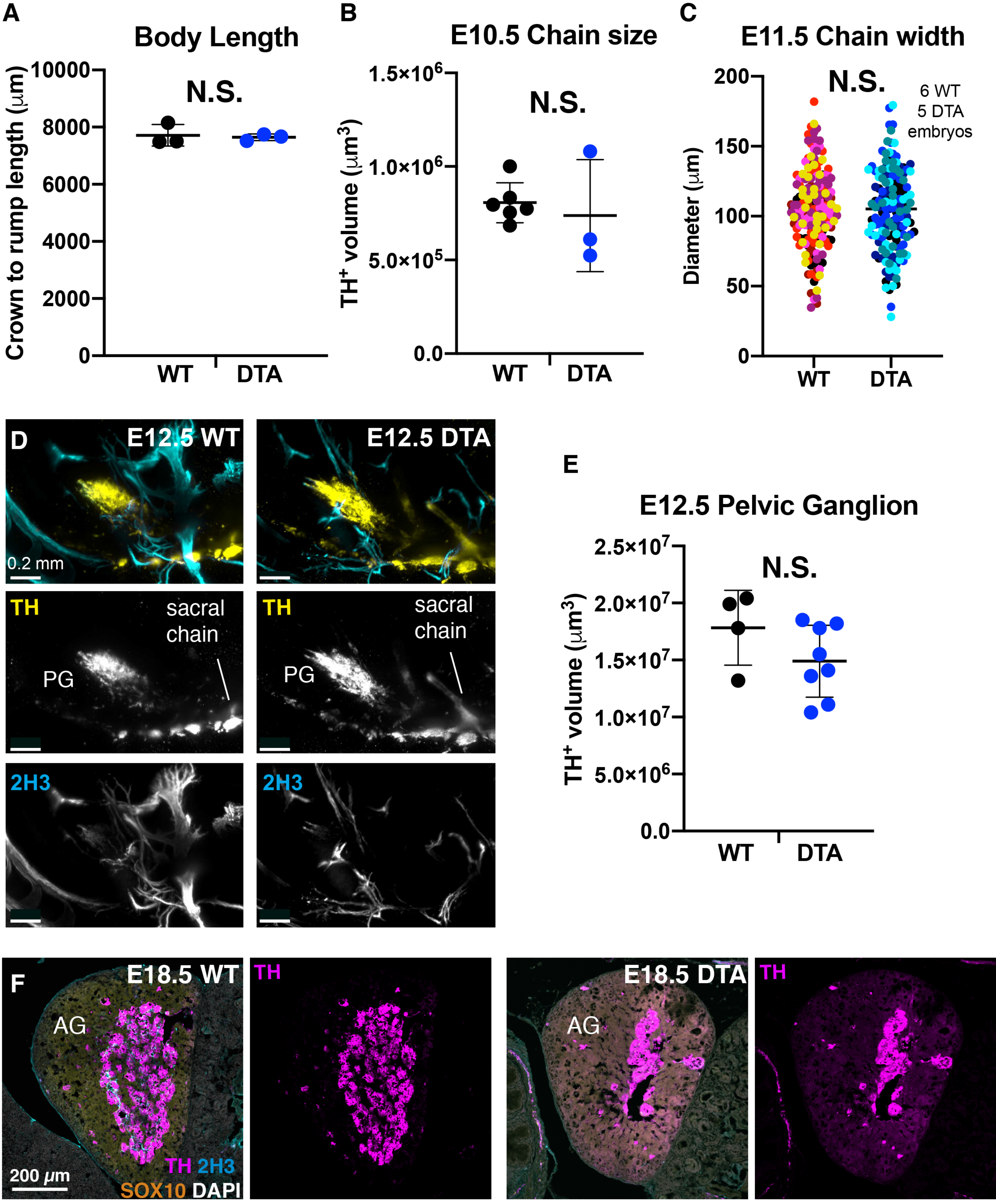


Supplementary Figure 2

**Supplementary Figure 2: Sympathoadrenal development in *Olig2*$^{Cre}$*; R26R*$^{DTA}$ embryos. (A)** Body length of the same E12.5 *Olig2*$^{Cre}$*; R26R*$^{DTA}$ and corresponding littermate control embryos. Error bars represent standard deviation and the middle bar represents mean. **(B)** Volume of $TH^+$ domains in E10.5 embryos. **(C)** Violin plots showing the distribution of the diameter (long axis) of sympathetic chain ganglia on transversal sections through E11.5 embryonic trunks at brachial and lumbar levels. Each datapoint represents one raw measurement. The six colors of datapoints represent the five mutants and six control animals analyzed for the experiment. **(D)** Pelvic ganglia in the E12.5 WT and *Olig2*$^{Cre}$*; R26R*$^{DTA}$ (accompanying quantification of volume shown on the right panel, each point is one ganglion). Scale bar = 0.2 millimeters. **(E)** Quantification of the distance from the sympathetic ganglia to the distal tip of the outgrowing sympathetic nerve in the forelimb. For all volumetric analysis shown in this figure, each datapoint represents the average volume of both left/right sympathetic ganglia volumes from an individual embryo except for (F), where ganglia were calculated as separate measurements. Student's t-test was used to determine statistical significant differences between the lengths and volumes of the sympathetic ganglia of *Olig2*$^{Cre}$*; R26R*$^{DTA}$ versus control littermates (**$p < 0.005$; *$p < 0.05$; N.S. no significant difference detected). The test reveals significant differences in posterior chain and adrenal/paraganglia volumes between normal and motor-ablated conditions at E12.5, whether left/right ganglia are averaged per embryo (as shown) or calculated as individual datapoints (not shown). **(F)** Adrenal glands from E18.5 *Olig2*$^{Cre}$*; R26R*$^{DTA}$ embryos and wild-type littermates immunostained for chromaffin cells using TH antibody.

### Sympathetic nervous system in alternate genetic model of motor ablation

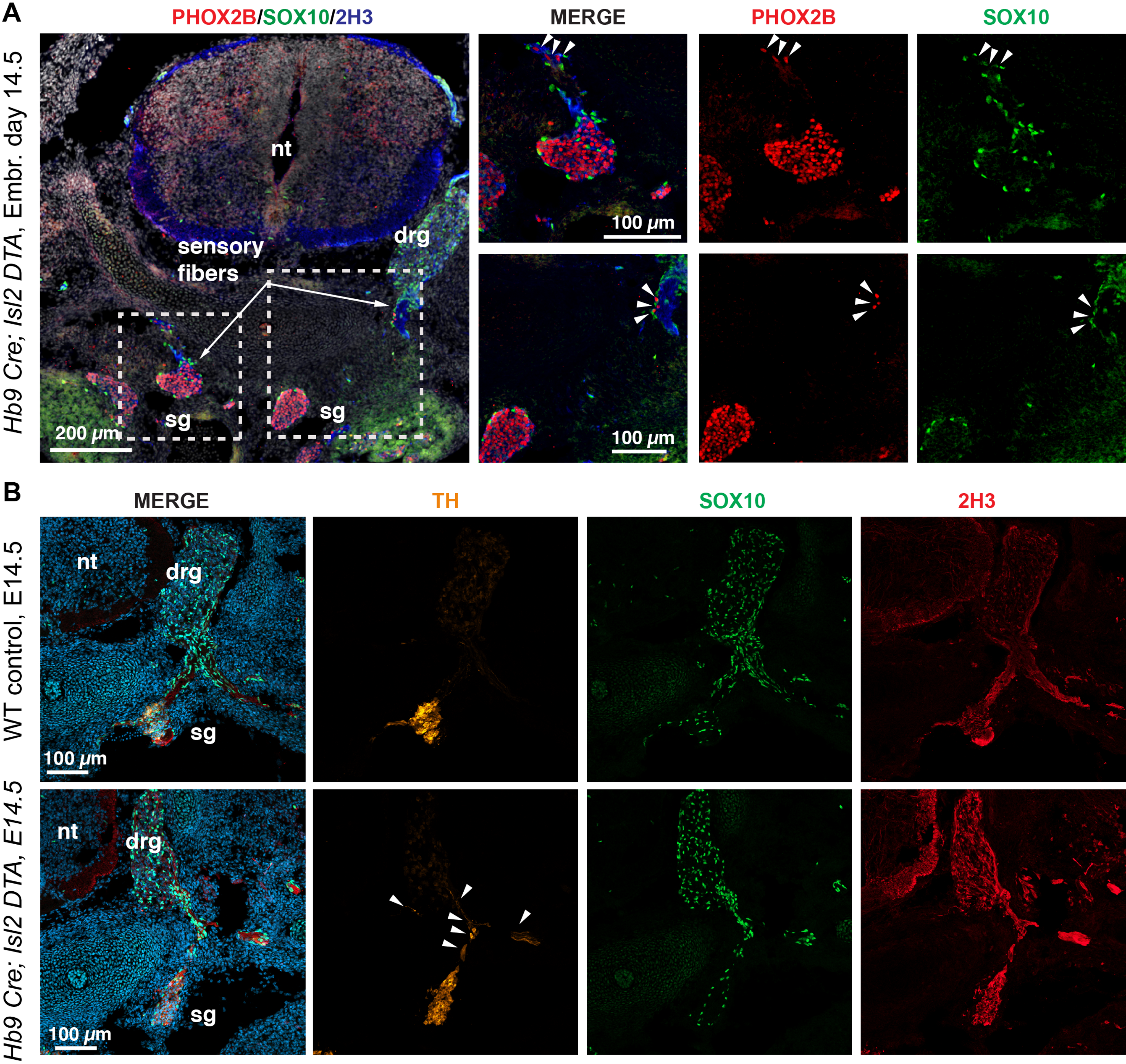


**Supplementary Figure 3**

**Supplementary figure 3: Fragmentation of sympathetic ganglia recapitulated in a second genetic model of motor nerve ablation. (A)** Immunostaining for PHOX2B, SOX10, and 2H3, and **(B)** immunostaining for TH, SOX10, and 2H3. All images show transverse sections through the trunk of $Hb9^{Cre}$:$Isl2^{DTA}$ embryo at E14.5. Left, overview merged image. Arrowheads point to SOX10/PHOX2B double-positive cells or SOX10/TH double-positive cells associated with the sensory nerves, positioned at a distance from the main sympathetic ganglia. Scale bars in overview in **(A)** = 200 micrometers, all other scale bars = 100 micrometers. NT: neural tube, DRG: dorsal root ganglion, SG: sympathetic chain.

Sensory nerve phenotype in motor-ablated embryos

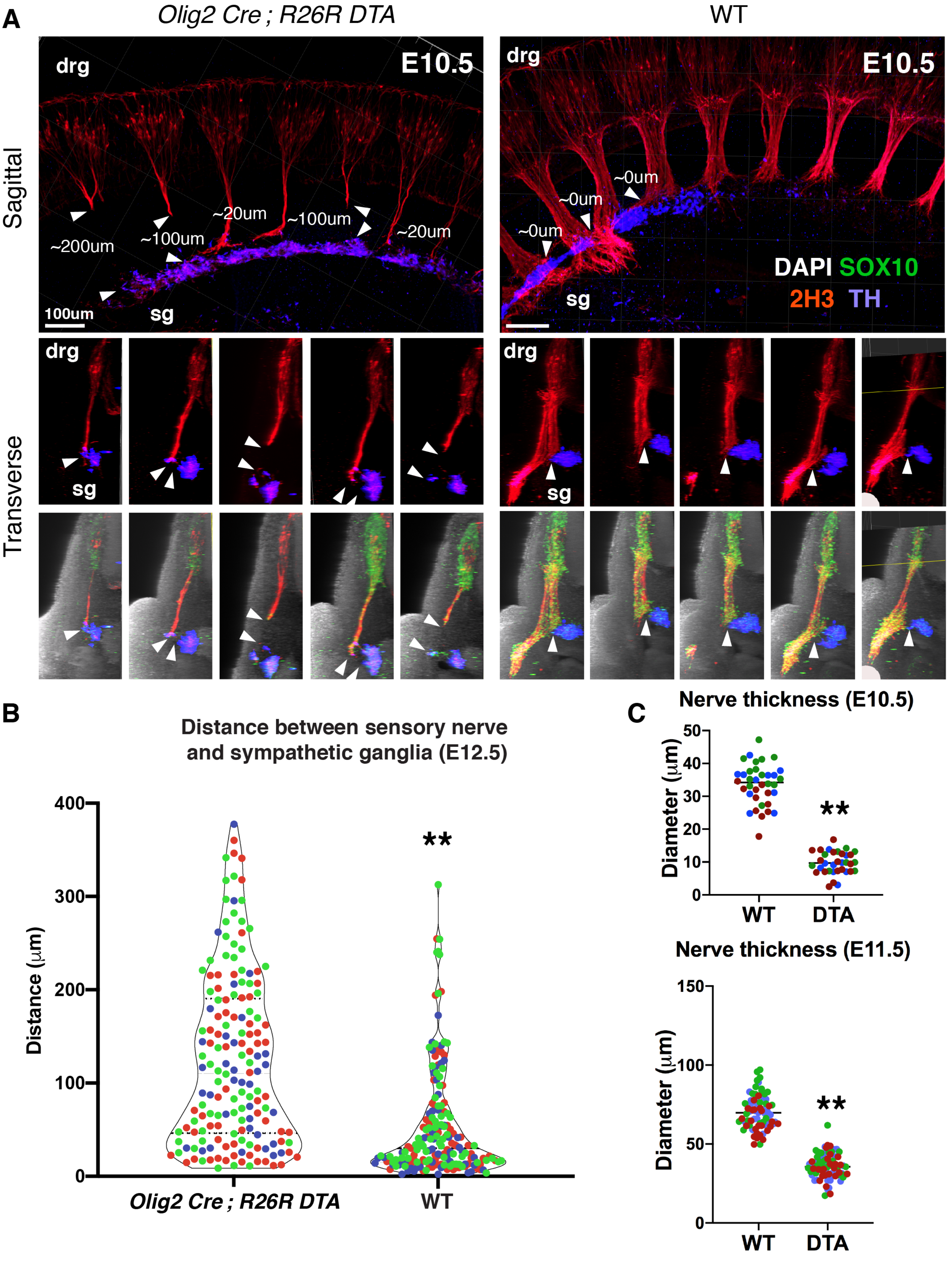


Supplementary Figure 4

**Supplementary figure 4: Defective sensory axon growth following motor nerve ablation.** **(A)** Sagittal (top) and transversal views (bottom panels) of E10.5 *Olig2*$^{Cre}$*;R26R*$^{DTA}$ embryo or WT littermate stained for 2H3, SOX10 and TH. The embryos are the same shown in Figure 3A. Each transversal view corresponds to a different section of the sympathetic chain anlagen and associated peripheral innervation. Arrowheads indicate the distance between nerve tip and sympathetic anlagen. Scale bar = 100 micrometers. **(B)** Quantifications of distances between TH+ sympathetic ganglia and 2H3+ / TH- nerve fibers. The analyzed images were from transversal sections of E12.5 *Olig2*$^{Cre}$*;R26R*$^{DTA}$ and WT littermate embryos stained with 2H3, SOX10 and TH. Each color represents the distribution of measurements across the body axis for 3 different embryos per genotype. Students t-test was used to determine statistically significant differences in the average distance between sympathetic ganglia and ventral root, comparing between transgenic and control embryos (** p value < 0.005). drg: Dorsal root ganglia, sg: sympathetic ganglia. **(C)** Diameter of peripheral nerves in motor-ablated and control trunks, measured in whole mount at E10.5 and E11.5, using manual distance measurement with Imaris software.

### Spatial analysis of nerve association of neural crest cells at E10.5

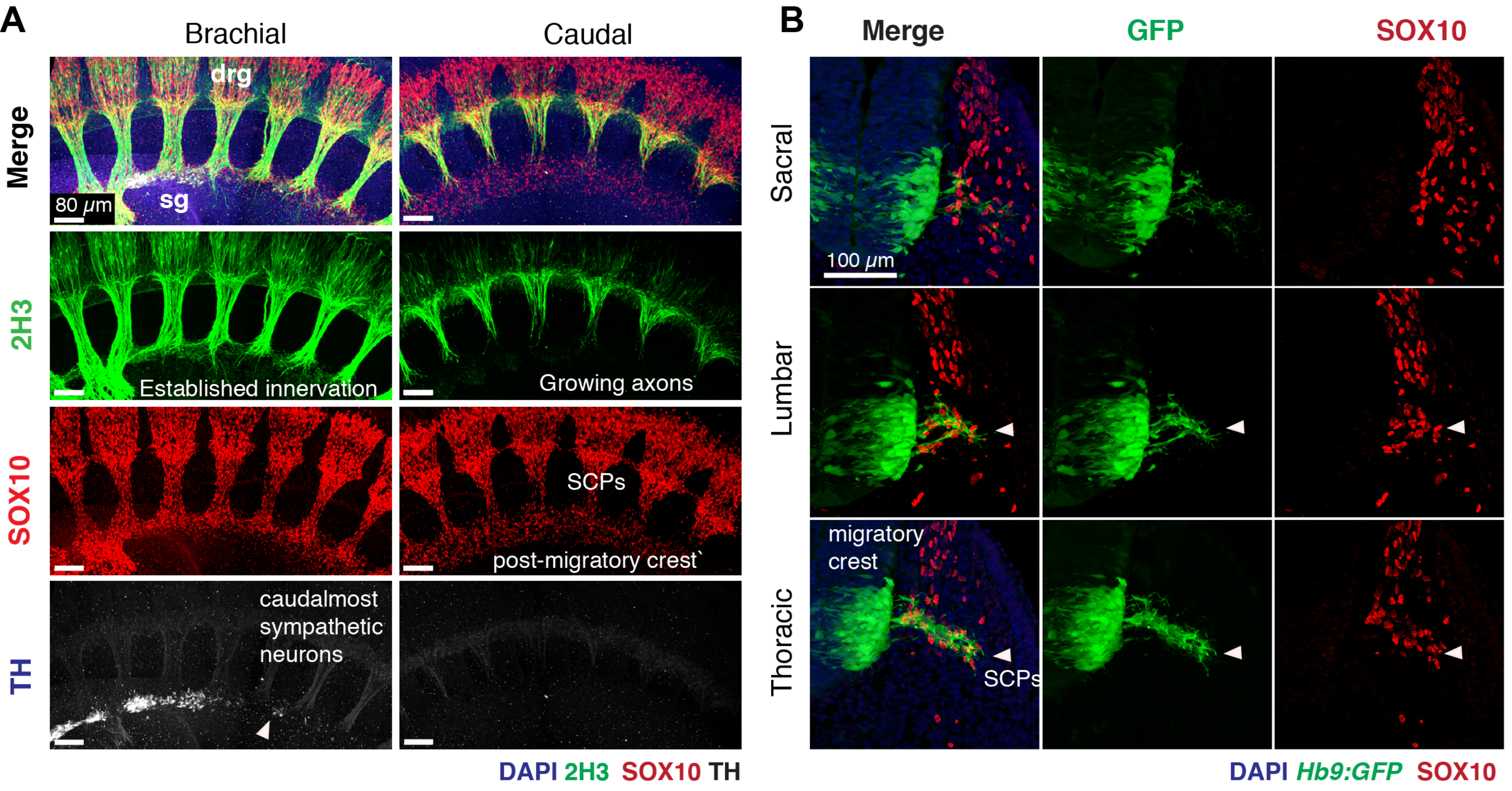


### Lineage tracing *Plp1*-expressing sympathetic progenitors from E10.5

*Plp1*$^{CreERT2}$ ; *R26R* $^{YFP}$, induced with 1mg tamoxifen at E10.5, harvest at E13.5

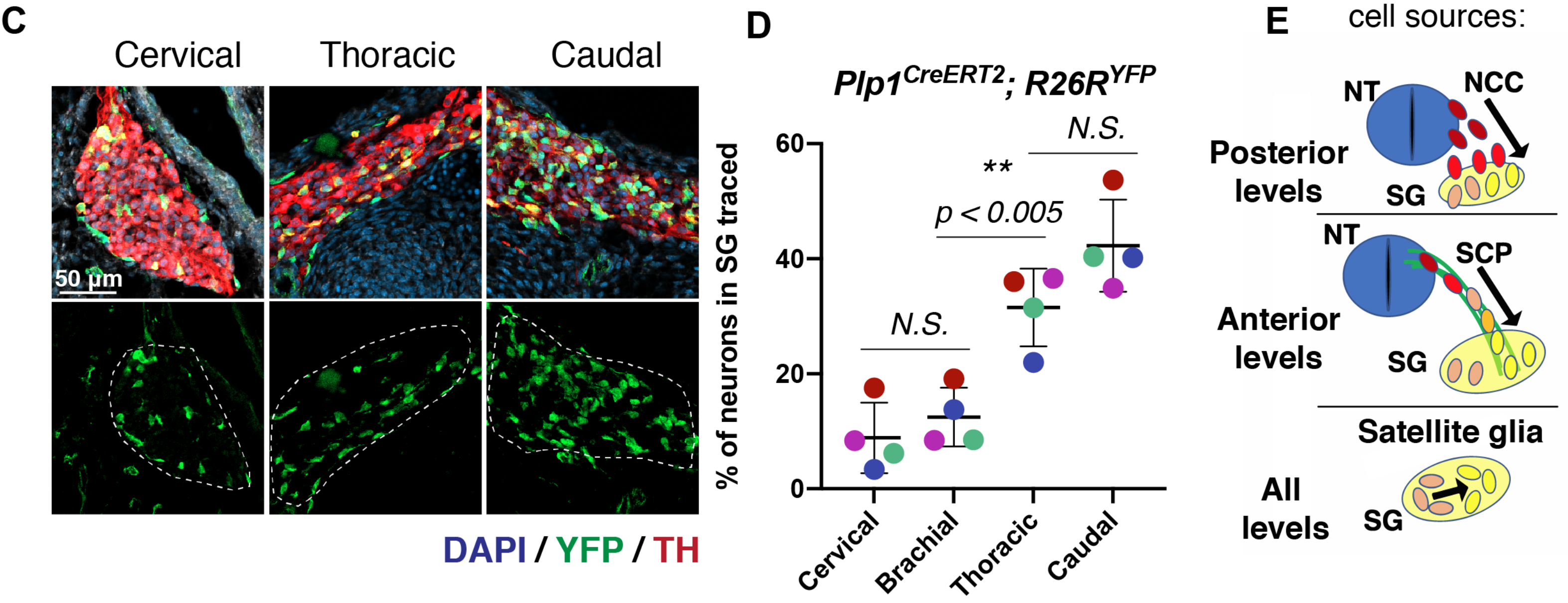


Supplementary Figure 5

**Supplementary Figure 5: Motor nerve outgrowth regulates the switch from free to nerve-mediated neural crest migration. (A)** Sagittal view of 10.5 embryo wholemount stained for peripheral innervation (2H3), SOX10, and the sympathetic anlagen (TH). At brachial levels (left panels) peripheral innervation is already well established and the sympathetic anlagen is in contact with sensorimotor fibers, whereas more caudally (right panels), sensorimotor axons have yet reached the sympathetic anlagen. Note that no $TH^+$ cells exist yet in the presumptive autonomic domain in caudal (bottom right) while sympathetic maturation has already begun at brachial level. Scale bar = 80 micrometers. The embryo shown is the same as shown in Figure 3A. The images are representative of at least 3 control embryos. **(B)** Immunostained transversal sections of an E10.5 *Hb9:GFP* embryo at increasingly anterior levels through the trunk to create a pseudo-timeseries. First (exemplified by the sacral level, top panels) neural crest cells are ventrally migrating freely and motor nerves have just begun to extend from the ventral neural tube. Next (exemplified by lumbar level, middle panels) motor nerves grow ventral-laterally to intersect with the path of neural crest cells, some of which begin to associate (shown with arrowheads). The flow of free NCC migration stopping is reflected in the sharp decrease in cell density ventral to the motor nerves. Finally (exemplified by thoracic levels, lower panels) nearly all NCC that reach motor nerves become associated with them (no cells freely migrating NCC are visible ventral to the motor root), marking the transition from free- to nerve-associated cell migration. Scale bar = 100 micrometers. **(C)** Immunostaining for TH and YFP on sagittal sections through sympathetic ganglia at different anatomical locations of *Plp1$^{CreERT2}$; R26R$^{YFP}$* embryos. Embryos were injected with 1mg tamoxifen at E10.5 and analyzed at E13.5. Scale bar = 50 micrometers. **(D)** Quantifications of TH/YFP double-positive cells traced in each type of ganglia. Each symbol corresponds to a different embryo. Student's t-test was used to determine statistically significant levels of tracing among different anatomical locations (**p value < 0.005; N.S., no significant difference detected). SG: sympathetic ganglia. **(E)** Schematic showing the various cell sources for sympathetic chain neurons.

### Cervical ganglia morphology following cholinergic cell ablation

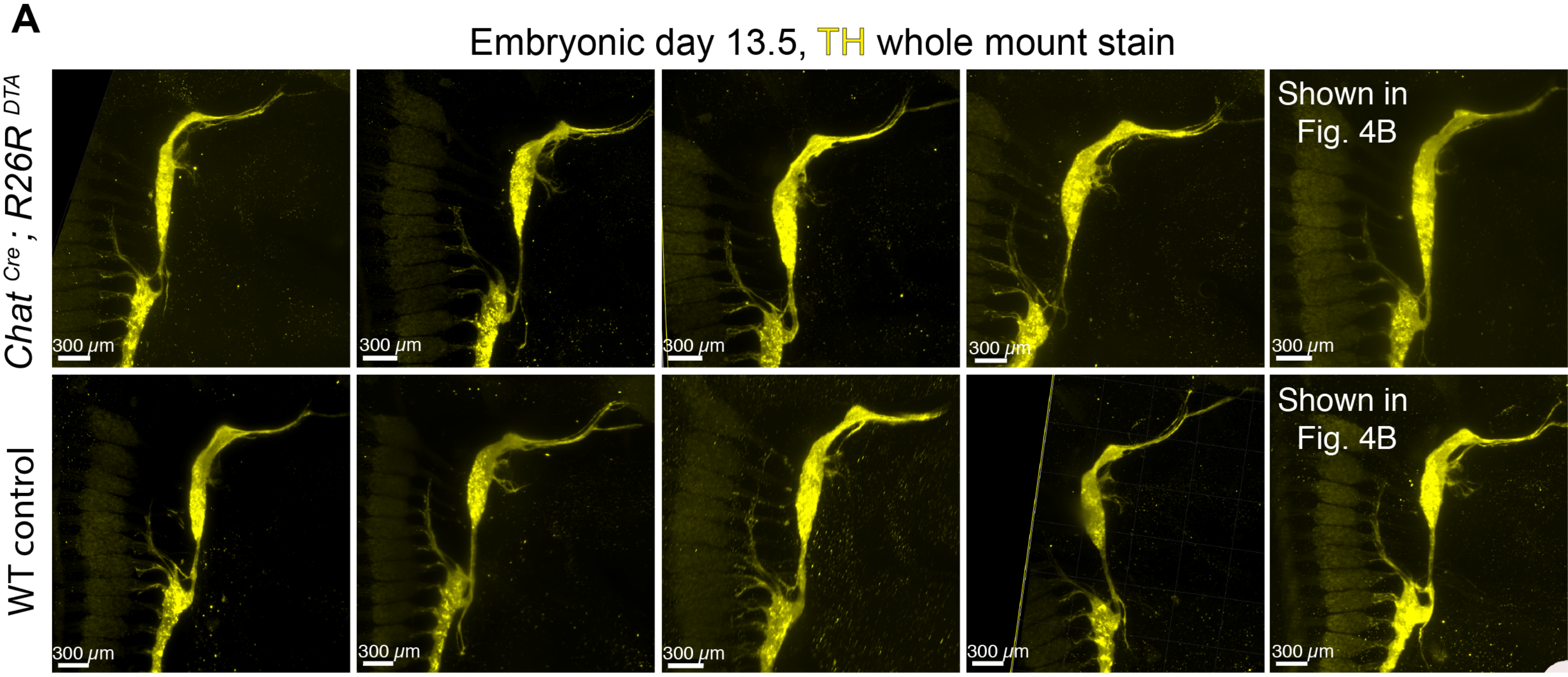


### Lineage tracing of *Chat*-expressing cells in the neural tube and peripheral ganglia

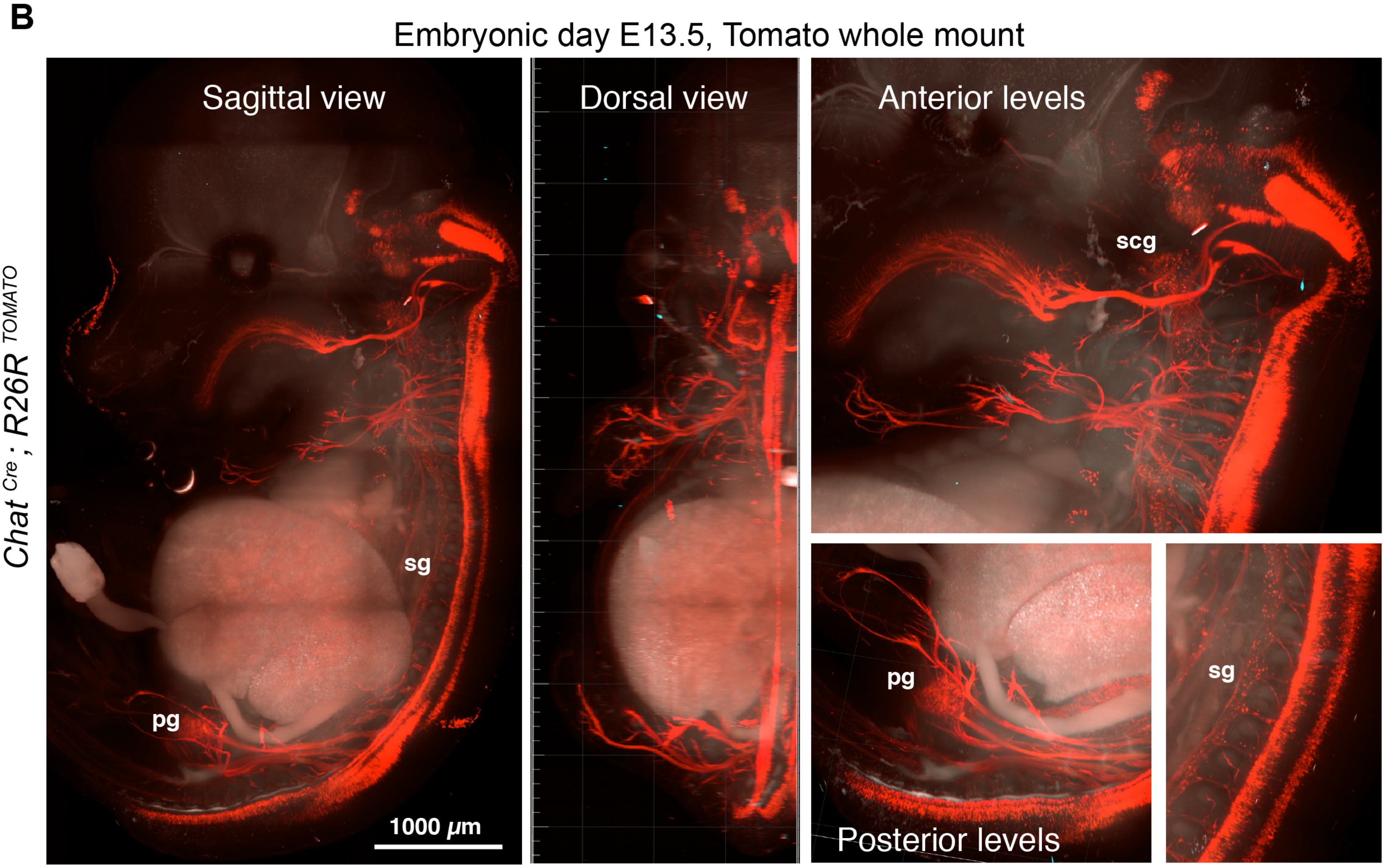


**Supplementary Figure 6**

**Supplementary Figure 6: Ablation of mature cholinergic neurons does not appear to have a strong impact on early cervical sympathetic ganglia morphogenesis. (A)** Examples of cervical ganglia (sagittal view) from five $Chat^{Cre};R26R^{DTA}$ embryos and five WT controls at E13.5. Scale bars = 300 micrometers. **(B)** Direct fluorescence from a $Chat^{Cre};R26R^{TOMATO}$ embryo, viewed from sagittal (left) and dorsal (middle) angles. Right panels show increased magnification of cervical, sacral, and thoracic regions. Scale bar = 1000 micrometers. scg: Superior cervical ganglion, pg: pelvic ganglion, sg: sympathetic ganglia.

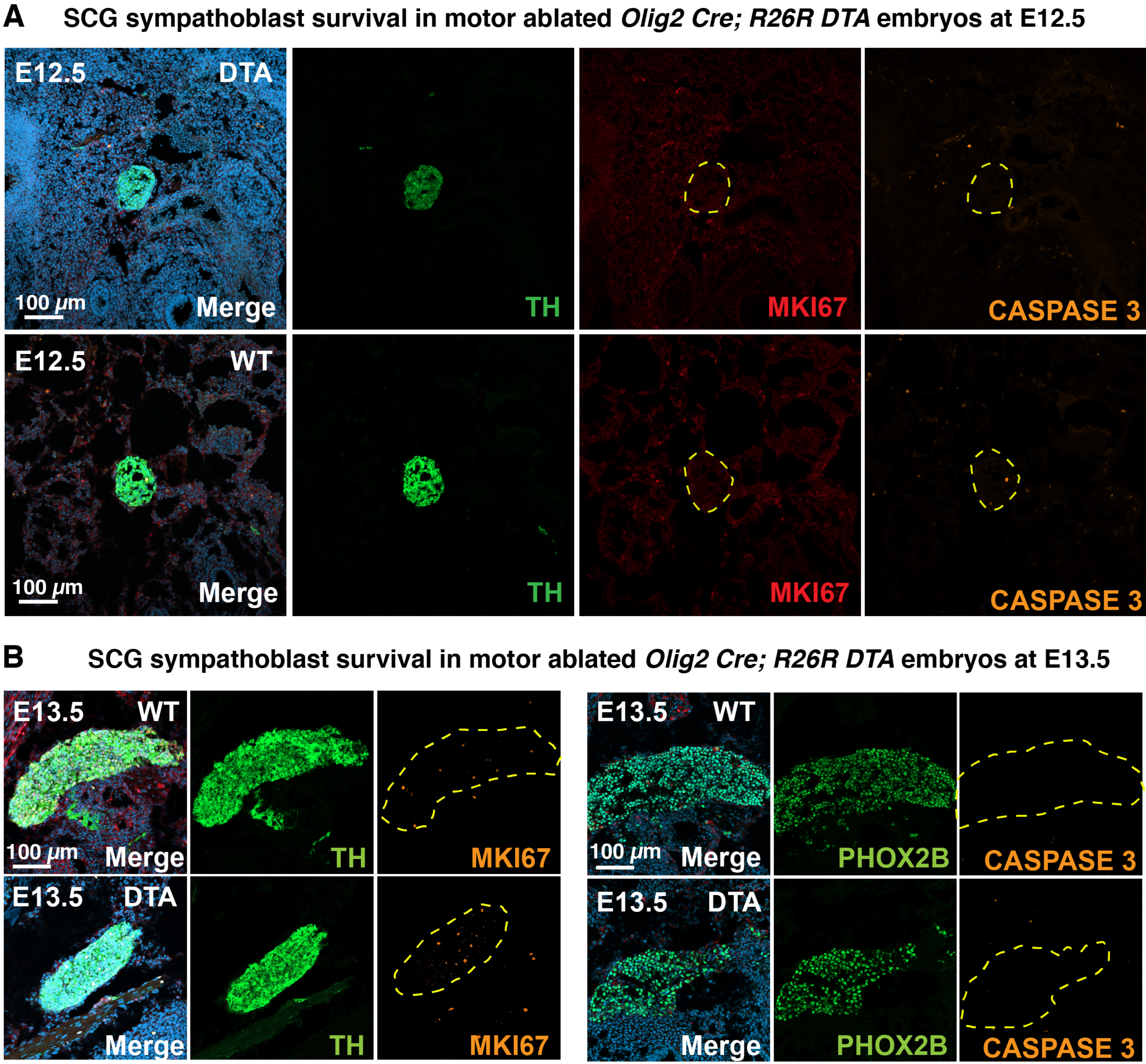


Supplementary Figure 7

**Supplementary Figure 7: Cervical sympathoblast proliferation and survival appear unaffected by motor neuron ablation during morphogenesis. (A)** Immunostaining for TH, proliferating cells (MKI67) and apoptotic cells (Cleaved caspase 3) on transverse sections through superior cervical ganglia of *Olig2$^{Cre}$; R26R$^{DTA}$* (top panels) and control littermate embryos (bottom panels) at E12.5. Scale bar = 100 micrometer. Images shown are representative of N=3 experimental and 3 control embryos. **(B)** Immunostaining for autonomic markers (TH left panels, PHOX2B right panels), MKI67 (left panels) and Cleaved caspase 3 (right panels) on sagittal sections through superior cervical ganglia of *Olig2$^{Cre}$; R26R$^{DTA}$* and control littermate embryos at E13.5. Scale bar = 100 micrometers. Images shown in (B) are representative of N = 1 embryo per genotype. Dotted lines represent outline of ganglia. Proliferating and apoptotic cells are scattered thinly throughout the tissue and very rarely observed in the ganglia, for either genotype.

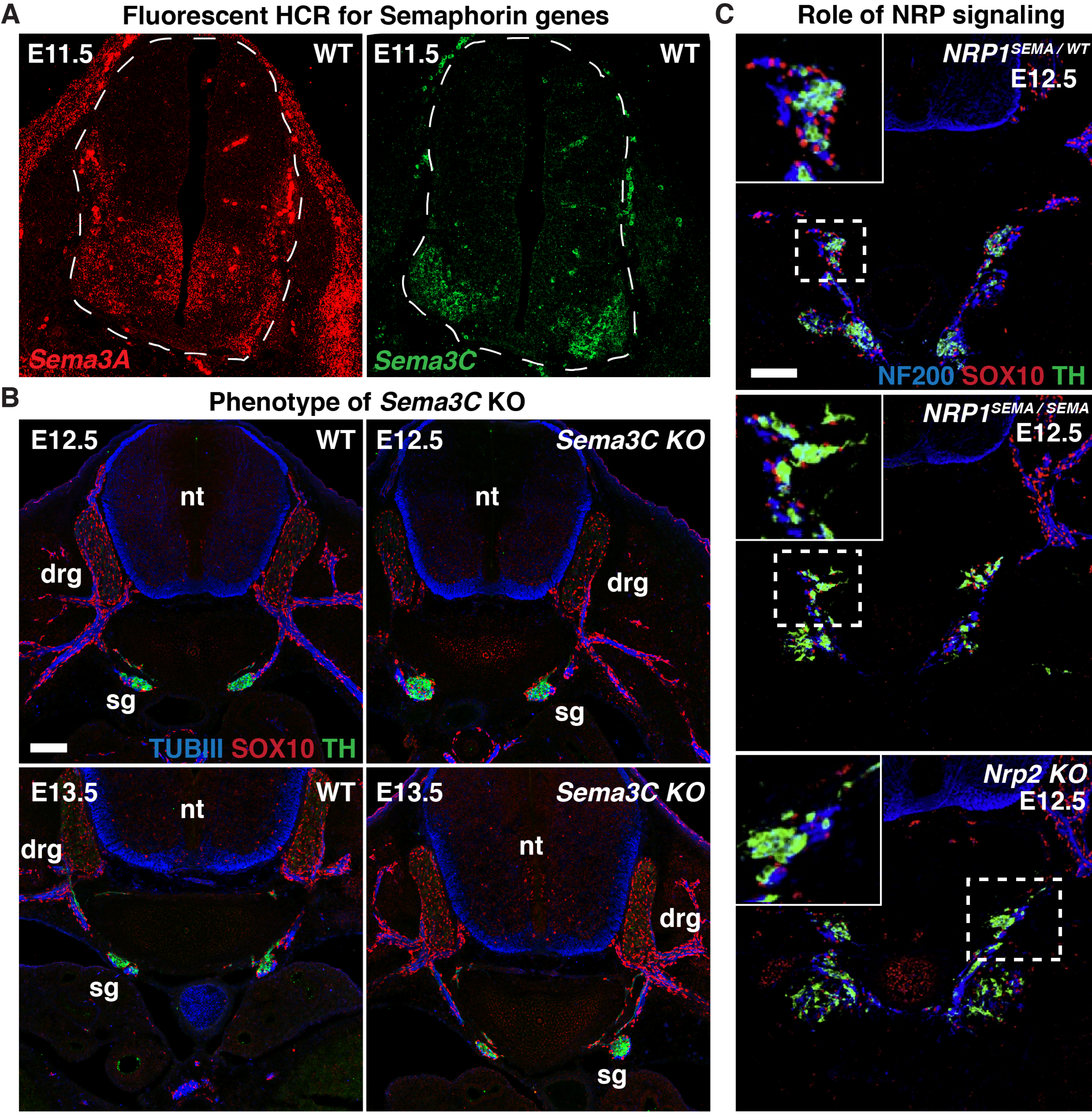


**Supplementary Figure 8**

**Supplementary Figure 8: Sema3C does not affect sympathoblast positioning. (A)** In situ hybridization chain reaction (HCR) of Semaphorin3A (top panel) and Semaphorin 3C (bottom panel) in transverse sections of embryonic day 11.5 mouse embryonic trunk (neural tube outlined with the dotted line). DRG = Dorsal root ganglia. MN = motor neuron. dNT = dorsal neural tube. **(B)** Sema3C KO embryos does not display a phenotype in sympathetic nervous system assembly and are similar to littermate control at both E12.5 (left panels) and E13.5 (right panels). Immunofluorescence staining of TH (green), Sox10 (Red), and TUBIII (blue) to mark sympathetic neurons, glial, and peripheral nerves respectively. **(C)** Phenotype in the sympathetic ganglia of $Nrp1^{SEMA/+}$ (top), $Nrp1^{SEMA/SEMA}$ (middle), and $Nrp2^{-/-}$ (bottom) embryos. Immunofluorescence staining of TH (green), Sox10 (Red), and NF200 (blue) to mark sympathetic neurons, glial, and peripheral nerves respectively.
